# Linking prokaryotic genome size variation to metabolic potential and environment

**DOI:** 10.1101/2022.10.20.512849

**Authors:** Alejandro Rodríguez-Gijón, Moritz Buck, Anders F. Andersson, Dandan Izabel-Shen, Francisco J. A. Nascimento, Sarahi L. Garcia

## Abstract

While theories and models have appeared to explain genome size as a result of evolutionary processes, little work has shown that genome sizes carry ecological signatures. Our work delves into the ecological implications of microbial genome size variation in benthic and pelagic habitats across environmental gradients of the brackish Baltic Sea. While depth is significantly associated with genome size in benthic and pelagic brackish metagenomes, salinity is only correlated to genome size in benthic metagenomes. Overall, we confirm that prokaryotic genome sizes in Baltic sediments (3.47 Mbp) are significantly bigger than in the water column (2.96 Mbp). While benthic genomes have a higher number of functions than pelagic genomes, the smallest genomes coded for a higher number of module steps per Mbp for most of the functions irrespective of their environment. Some examples of this functions are amino acid metabolism and central carbohydrate metabolism. However, we observed that nitrogen metabolism was almost absent in pelagic genomes and was mostly present in benthic genomes. Finally, we also show that Bacteria inhabiting Baltic sediments and water column not only differ in taxonomy, but also in their metabolic potential, such as the Wood-Ljungdahl pathway or the presence of different hydrogenases. Our work shows how microbial genome size is linked to abiotic factors in the environment, metabolic potential and taxonomic identity of Bacteria and Archaea within aquatic ecosystems.

## INTRODUCTION

Genomes in Bacteria and Archaea are information-rich (1), and known to range in size from 0.1 to 16 million base pairs (Mbp) (2). They can vary over evolutionary time through genomic expansions and contractions via genetic drift, selection, homologous recombination, deletions and insertions (3–9). Moreover, evolutionary studies have revealed extremely rapid and highly variable flux of genes (10) with evolutionary forces acting on individual genes (5). With all these evolutionary forces acting on the genes, we can presume that gene content and, by consequence, genome size has an ecological meaning. Indeed, genome size has been linked to phylogenetic history (11,12), lifestyle such as free-living, particle attached or host-associated (4,13,14), or environment such as marine, freshwater, different types of sediments or different hosts in host-associated microorganisms (2,15,16). We aim to delve into the ecological implications of genome size in aquatic microorganisms, with emphasis on metabolic potential using a brackish environment as a model.

In the last decade, aquatic microorganisms have been extensively sampled and now have a large representation in genomic and metagenomic datasets (17). Their genome size spans from 0.5 to 15 Mbp with an average of 3.1 Mbp (2). Aquatic environments are heterogeneous and many different abiotic factors, such as salinity and depth, could be linked to microbial genome size variation. For example, pelagic microbes inhabiting deep marine environments are estimated to present bigger genome sizes than those in shallow marine waters (18,19). Within freshwater ecosystems, isolates from the family Methylophilaceae (class Gammaproteobacteria) show a smaller genome size for pelagic than for sediment dwellers (20). Additionally, two studies have shown that marine Cyanobacteria have smaller genome sizes than freshwater (21,22). This literature already provides some insights on how genome size is linked to the environmental heterogeneity of freshwater and marine ecosystems. Yet, the studies are limited either to a specific marine station, or specific microbial lineages and it remains a question if these findings are applicable more widely. Moreover, genome size variation in the brackish realm remains debated: Actinobacteria in the brackish Baltic Sea show bigger genome sizes than in freshwaters (23), while picocyanobacteria show the opposite trend (22). Additionally, further research must be done to elucidate the link between genome size and abiotic factors within aquatic environments, particularly brackish water bodies.

In the link between abiotic factors in the environment and genome size, gene content is selected accordingly. Metagenomic studies show that a vast majority of the genes in Bacteria and Archaea are specific to particular environments, whereas very few genes are being shared between environments (24). This remarks how relevant is the relationship between niche specificity and lineage specific functional traits (25). These functional capabilities also differ between water column and sediments in Bacteria and Archaea in both marine (26,27) and freshwater environments (28). Since gene repertoires and genome size are related, they must be considered together with environmental gradients to better understand niche-specificity and the ecology of different prokaryotic lineages.

In this research article, we provide a comprehensive analysis to show the ecological implications of genome size variation of Bacteria and Archaea in pelagic and benthic communities in the Baltic Sea. Specifically, we investigate; i) how genome size varies across abiotic factors and taxonomic lineages of Bacteria and Archaea from sediments and water column, ii) what relationship can be found between genome size and the number of metabolic capabilities in Bacteria in the Baltic Sea, and iii) which taxa and metabolic pathways contrast between pelagic and benthic Bacteria. To achieve this, we selected 111 pelagic and 59 benthic metagenomic samples that were previously published, and we provide one new and unpublished benthic dataset with 49 metagenomic samples. We use these 219 metagenomes to study genome size distribution across sediments and water column in the Baltic Sea (**Figure 1** and **Table S1**). For this, we use two different approaches to study genome size: the estimated average genome size (AGS) per metagenomic sample, and the estimated genome size of bacterial and archaeal metagenome-assembled genomes (MAGs). Our results show that Bacteria and Archaea with larger genomes in the sediments present lower coding density than the smaller genomes in the water column. We also find that microorganisms inhabiting the Baltic benthos have more metabolically-versatile genomes than pelagic prokaryotes, which mean they code for a wider range of metabolic capabilities. Finally, we also find that functions involved in the nitrogen metabolism are disproportionately more detected in benthic bacteria in the Baltic Sea.

**Figure 1.**
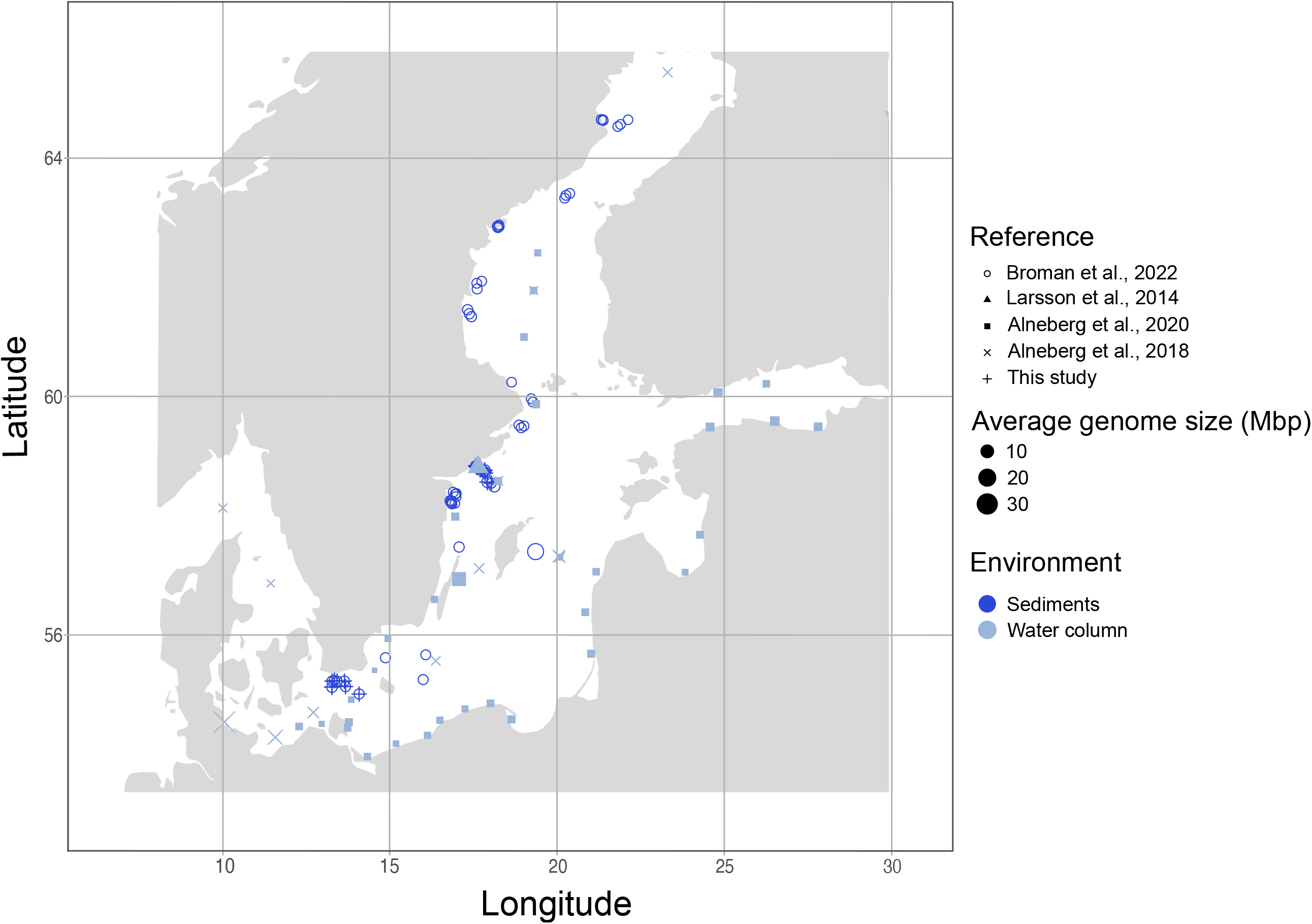
Overview of the sampling locations and average genome size (AGS) of metagenomes (108 from sediments in dark blue, and 111 from the water column in light blue). This figure shows the geographic location of all metagenomes used in this study. For exact coordinates see **Table S1**. Shape type indicates the reference and shape size indicates the AGS of the given metagenome.

## RESULTS AND DISCUSSION

### While depth is significantly associated with genome size in benthic and pelagic metagenomes, salinity is only correlated to genome size in benthic metagenomes

First, we calculated the average genome size (AGS) of metagenomes across the latitudinal gradient of the Baltic Sea comparing benthic and pelagic metagenomes. We observed that sediment-dwelling microbial communities present significantly larger AGS (mean = 6.01 Mbp, n = 108) than pelagic communities (mean AGS = 5.40 Mbp, n = 69) (Wilcoxon test, *p* < 0.01) (**Figure 2A**). We then evaluated the relationships between AGS of metagenomes and each of the environmental variables (depth, salinity, temperature, and oxygen concentration) independently and their interactions (ANOVA type II). The AGS in the pelagic metagenomes is significantly associated only with depth and shows a negative correlation (**Supplementary Material 1** and **Figure S1**). Previous marine analysis have found the opposite effect, bigger genome sizes in deeper areas (19). However, our pelagic analysis only covers 5 meters of depth and explains 19% of the genome size variation in pelagic metagenomes. AGS in sediment metagenomes is significantly associated with both salinity and depth and most of the interactions of these variables (**Supplementary Material 1**). Both depth and salinity have a weak positive correlation with AGS in the sediment samples (**Figure S1**). To our knowledge,his is the first time where a positive correlation between genome size and salinity is reported for brackish sediment microorganisms.

**Figure 2.**
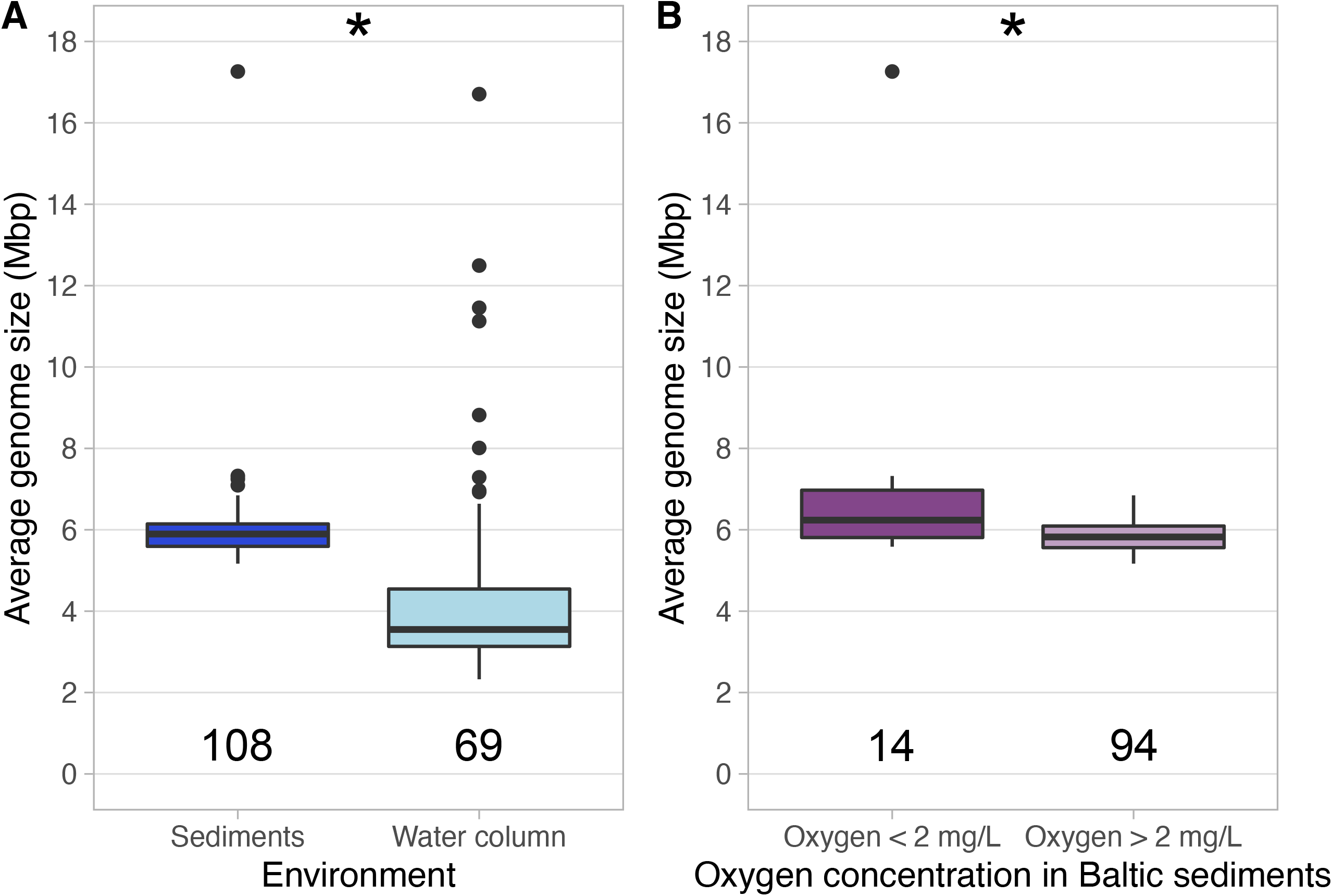
Boxplots showing the AGS distribution of Baltic metagenomes. **Panel A** indicates the AGS distribution in both water column and sediments. **Panel B** indicates the AGS distribution in metagenomes from sediments across the oxygen gradient (two groups, from 0 to 2 and from 2 to 12.45 mg/L). Stars in both panels indicate significant differences p < 0.05 (Wilcoxon non-parametric test).

Although the ANOVA did not show a significant effect of water oxygen concentration on genome size, we further investigate the effect of bottom water O2 concentration in the AGS of metagenomes from sediments. We separated these metagenomes into those from oxygen concentration 0 to 2 mg/L (mostly metagenomes from the dead zone) and those metagenomes with oxygen concentration 2 to 12.45 mg/L. We observe that benthic metagenomes from lower oxygen concentration (mean AGS = 7.08 Mbp, n = 14) present significantly bigger genome sizes than those in sediments with higher oxygen concentration (mean AGS = 5.85 Mbp, n = 94) (Wilcoxon test, *p* < 0.01) (**Figure 2B**). Complementarily, previous results show that the dead zone bacteria tend to be more metabolically similar to each other when compared to bacteria from oxic sediments (29). Our observation confirms a recent study that observed obligate anaerobes from diverse environments present bigger genome sizes than microaerophilic microorganisms (30). Moreover, previous studies show a positive correlation between genome size and nutrient concentration (19,31,32). Dead zones in the Baltic Sea are characterized by anthropogenic eutrophication (33), which would also promote bigger genome sizes. Altogether, these results indicate that high nutrient concentration and low oxygen concentrations in Baltic Sea dead zones may select for prokaryotes with bigger genome sizes.

### While average benthic estimated genome size is bigger than pelagic, the biggest genomes in the Baltic Sea were found in the water column

To compare genome size between pelagic and benthic bacteria and archaea, we used metagenome-assembled genomes (MAGs; >75% completeness and <5% contamination) from our metagenome datasets. Our dataset compiles 216 MAGs from the sediments that dereplicated into 56 representative genomospecies (95% average nucleotide identity). Additionally, 1920 pelagic MAGs were dereplicated into 340 representative genomospecies. We observe that 12 phyla were detected in both sediments and water column, while 16 phyla were specific to either habitat. Seven phyla were found specific to the sediment (14 representative MAGs) and nine phyla were specific to the water column (29 representative MAGs) (**Table 1**). Interestingly, only one genomospecies representative was binned from both sediments and water column. This genome representative belongs to genus *Mycobacterium* (phylum Actinobacteriota, mOTU_124/pelagic and mOTU_027/sediments in **Table S2**), a genus that is not commonly found on brackish surface waters (34). However, this genus was found to be abundant in sediments in some regions of the Baltic Sea, especially in anoxic areas close to Landsort (35). These differences on taxonomical composition in microbial communities between water column and sediments altogether with latitudinal changes in microbial biodiversity in the Baltic Sea (34) show how heterogeneous is the microbial composition of brackish environments.

**Table 1.**
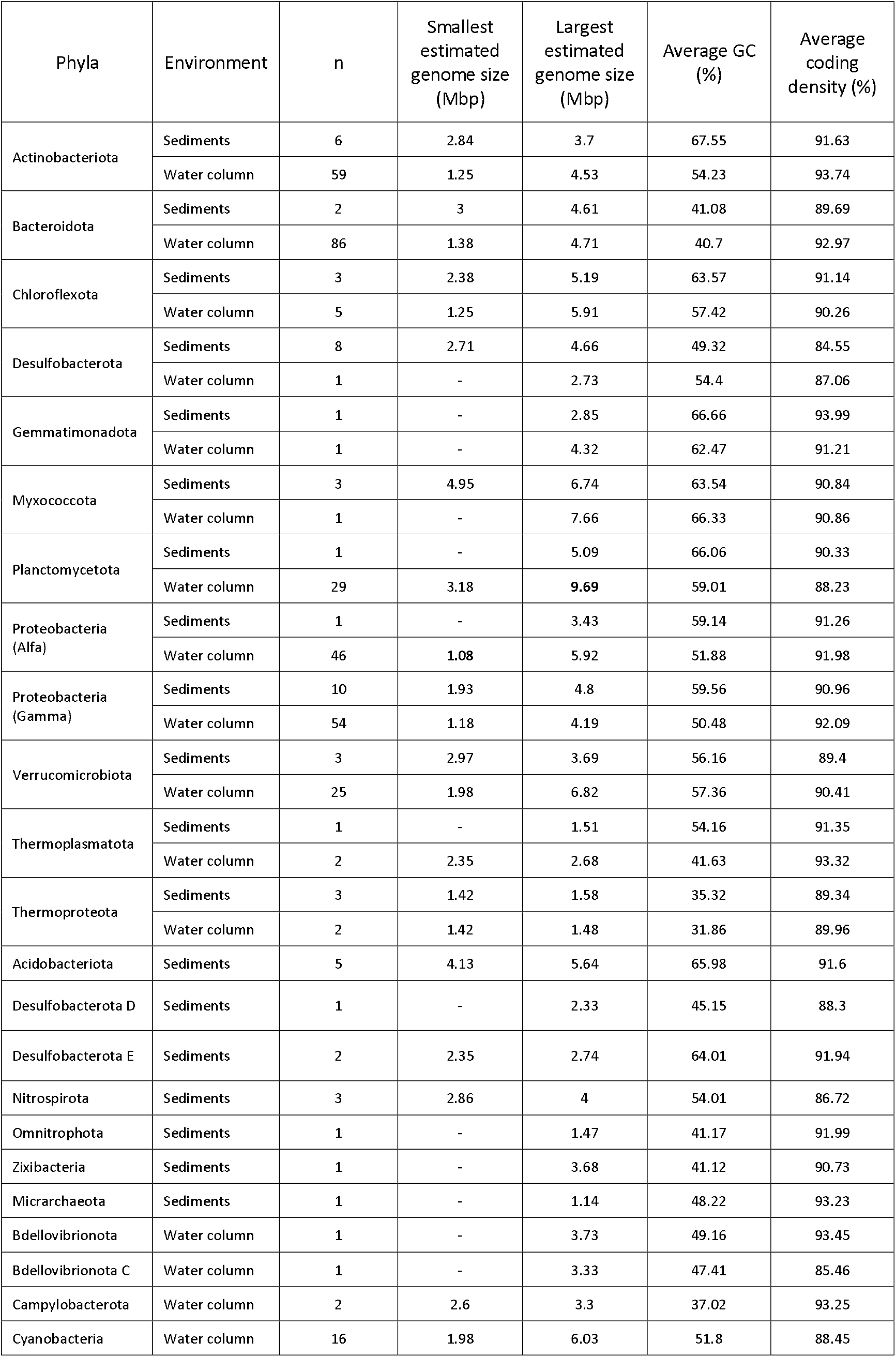
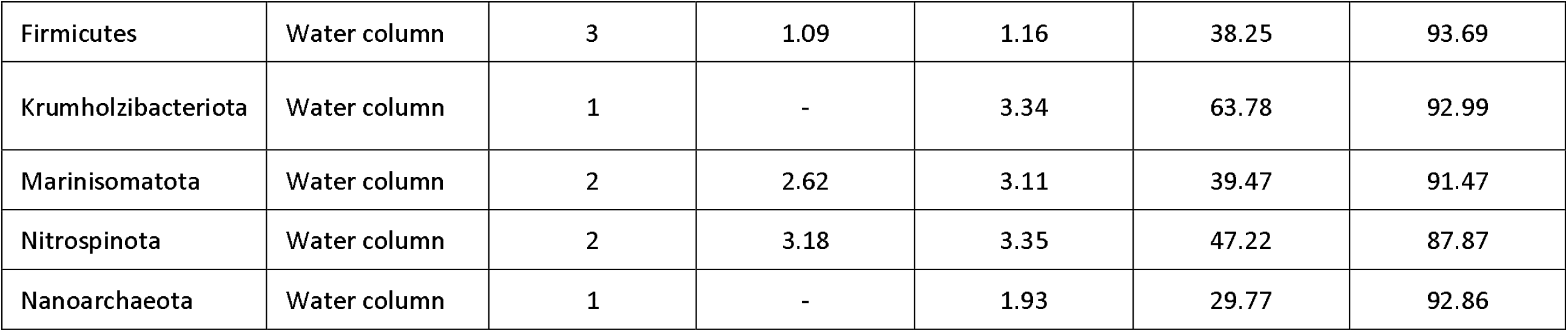
Summary of all 56 sediment and 340 water column representative MAGs (95% average nucleotide identity) with >75% completeness. Table includes phyla, environment (either water column or sediments), number of representative genomes (*n*), smallest and largest estimated genome sizes (Mbp) observed for each phylum, average GC content (%) and average coding density (%). When only one MAG is indicated, estimated genome size of that MAG is expressed in the sixth column.

Additionally, the eight largest representative genomes of the dataset were observed in the water column and belonged to phylum Planctomycetota (family Planctomycetaceae) (7.95 – 9.69 Mbp). It has been previously observed that Planctomycetota is the phylum containing the aquatic MAG with the biggest known estimated genome size (14.93 Mbp) (2). These large genomes contain large collections of genes that could be linked to the very complex cell structures and chromosomes observed in this phylum (36). Still, further research is necessary to understand if extant genome size in phylum Plactomycetota is the result of ecological adaptation to abiotic gradients. On the other side of the genome size spectrum, the representative MAG with the smallest estimated genome size belongs to class Alphaproteobacteria (family Pelagibacteraceae, 1.08 Mbp; **Table 1**). Bacteria from this family have been widely reported to be streamlined, abundant and ubiquitous across all salinity gradients (37,38).

Altogether, representative MAGs from sediments presented an average estimated genome size of 3.47 Mbp, which was significantly higher than for water column MAGs (2.96 Mbp) (*p* < 0.01) (**Figure 3A**). This was true also at the phyla, class, and order level (**Figure 3C** and **D**). Bacteria from sediments presented bigger estimated genome size on average (3.67 Mbp), followed by pelagic Bacteria (2.98 Mbp), pelagic Archaea (1.97 Mbp) and sediment Archaea (1.43 Mbp) (**Figure 3B**). This is supported by previous results, as Bacteria show bigger genome sizes than Archaea regardless of the environment (2). Moreover, the average estimated genome size of Baltic sediments (3.47 Mbp) is more similar to that of terrestrial microbial genomes (3.7 Mbp) (2). Additionally, genomes in the Baltic sediments have also lower coding density than pelagic (**Figure S2**). Our results corroborate previous findings that streamlining is common in pelagic marine environments (7,37,39,40) and pelagic brackish environments (23).

**Figure 3.**
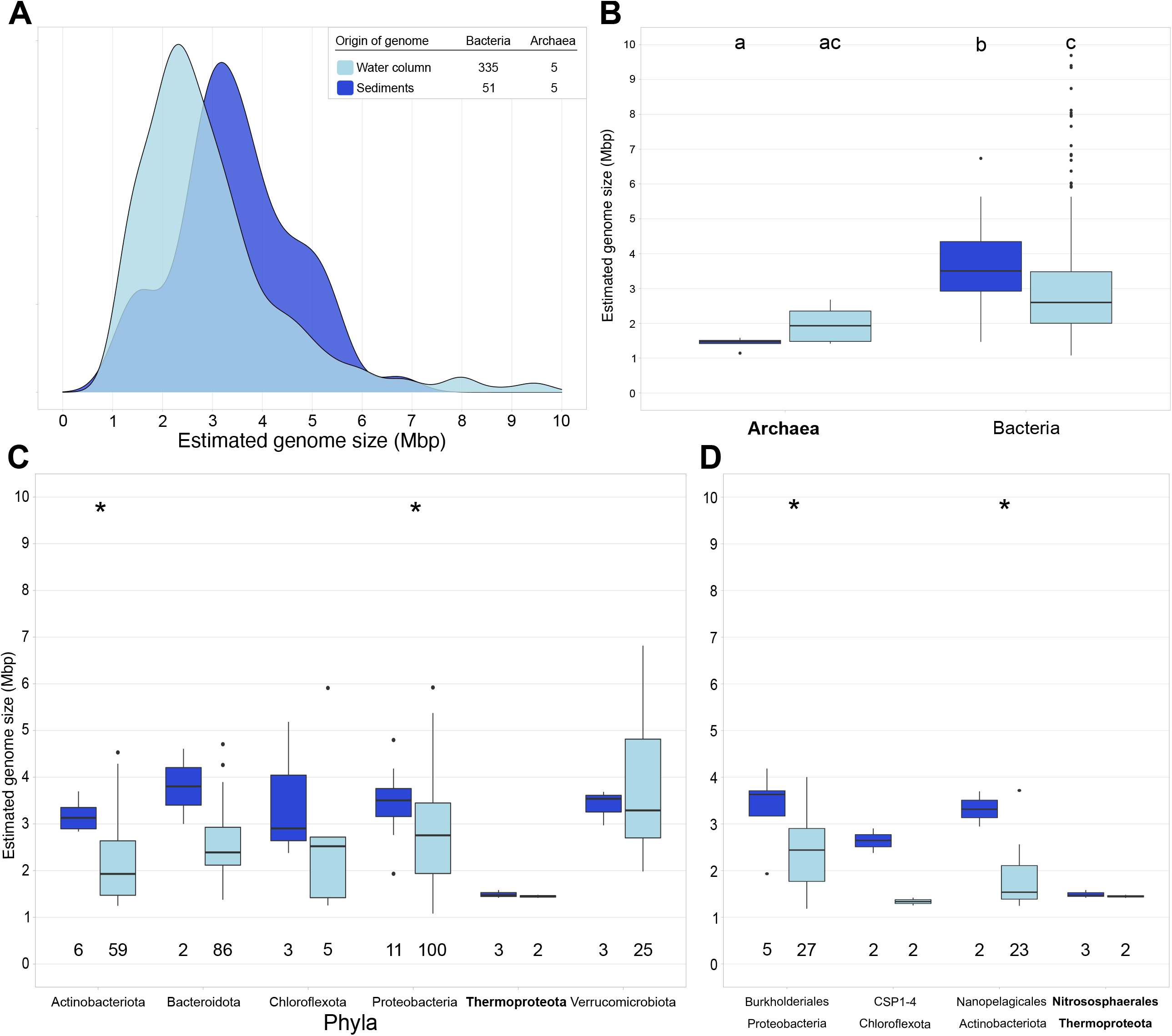
Overview of the estimated genome size in bacteria and archaea obtained from Baltic Sea sediments (dark blue) and water column (light blue) using only the 397 representative MAGs (95% average nucleotide identity) with >75% completeness. **Panel A** shows the genome size distribution of archaea and bacteria obtained from Baltic water column and sediments for a total of 397 representative genomes. **Panel B** shows the estimated genome size per domain and environment. Different letters indicate significant differences p < 0.05 (Kruskal-Wallis non-parametric test; multiple testing corrected with Benjamini-Hochberg). **Panel C** shows the estimated genome size per phylum. We selected only phyla with at least 2 MAGs in each environment. **Panel D** shows the estimated genome size per order. We selected only orders with at least 2 MAGs in each environment. Numbers below the boxes indicate the number of MAGs per environment. Stars in Panel C and D indicate significant differences p < 0.05 (Wilcoxon non-parametric test).

In our study, the average estimated AGS for the sediments (6.01 Mbp) and water column (4.44 Mbp) is larger than the average estimated genome size of the MAGs assembled and binned from the sediments (3.47 Mbp) and water column (2.96 Mbp), respectively (**Figure 2A** and **3A**). We calculated the AGS to estimate the average genome size of the whole microbial community. We used MicrobeCensus as a robust and accurate tool to calculate AGS (41). However, this AGS of metagenomes overestimates the genome size because of the viral and eukaryotic reads that might be present in the sample (42,43). On the other hand, there are two biases with looking at the average estimated genome size of MAGs. One, assembly and binning biases make a MAG an average 3.7% smaller than genomes from isolates in the same genomospecies (2). Second, a bias of overlooking all those bacteria and archaea that are hard to assemble and bin due to high genomic intrapopulation diversity (44). For these reasons, in our study, we use two methods that have different biases to answer the same question; how is the genome size of microorganisms distributed in the Baltic Sea. Irrespective of the method used, the average estimated genome size in sediments is larger than in the water column.

The available metadata for benthic and pelagic carbon concentration was not comparable due to methods and metrics used for analysis. This made it not possible to look for a clear ecological link between bigger genome size in benthic zones and nutrient concentration (**Table S1**). Luckily a previous study has compiled information on organic carbon stocks in the Baltic Sea (45): by collecting information from many different studies and years, they show that in average the top 10 cm of sediments contain between 2 and 4 times more organic carbon per area than the water column in the Baltic Sea. More organic carbon available for bacteria and archaea would also pose a lower pressure on the genome to streamline. From the results of our study, we hypothesize that one of the reasons pelagic microbial genomes are smaller than benthic microbial genomes is the difference in organic carbon availability. Altogether, a positive correlation between genome size and nutrient concentration has been shown before (19,31,32).

### Brackish pelagic microorganisms tend to show smaller genome sizes than marine and freshwater

In further analysis of genome size variation across salinity gradients, we compared the average estimated genome size of pelagic Baltic Sea MAGs to previously published genome size estimations (2). This comparison includes all taxonomic groups found in three large MAG-datasets (17,46,47) (completeness >75%) that includes 4051 freshwater representative MAGs, 2118 marine representative MAGs and 340 pelagic representative MAGs from the Baltic Sea. We found that the average estimated genome size of the brackish pelagic MAGs (2.96 Mbp) is lower than in marine MAGs (3.10 Mbp) (Kruskal-Wallis test, *p* < 0.01). Furthermore, we observed the largest average estimated genome size in freshwater MAGs (3.48 Mbp) (Kruskal-Wallis test, *p* < 0.01) (**Figure 4A**). These observed differences in genome sizes across different pelagic environments together with the previously observed differential functions (48) suggests that genome size has a potential signature across aquatic environments.

**Figure 4.**
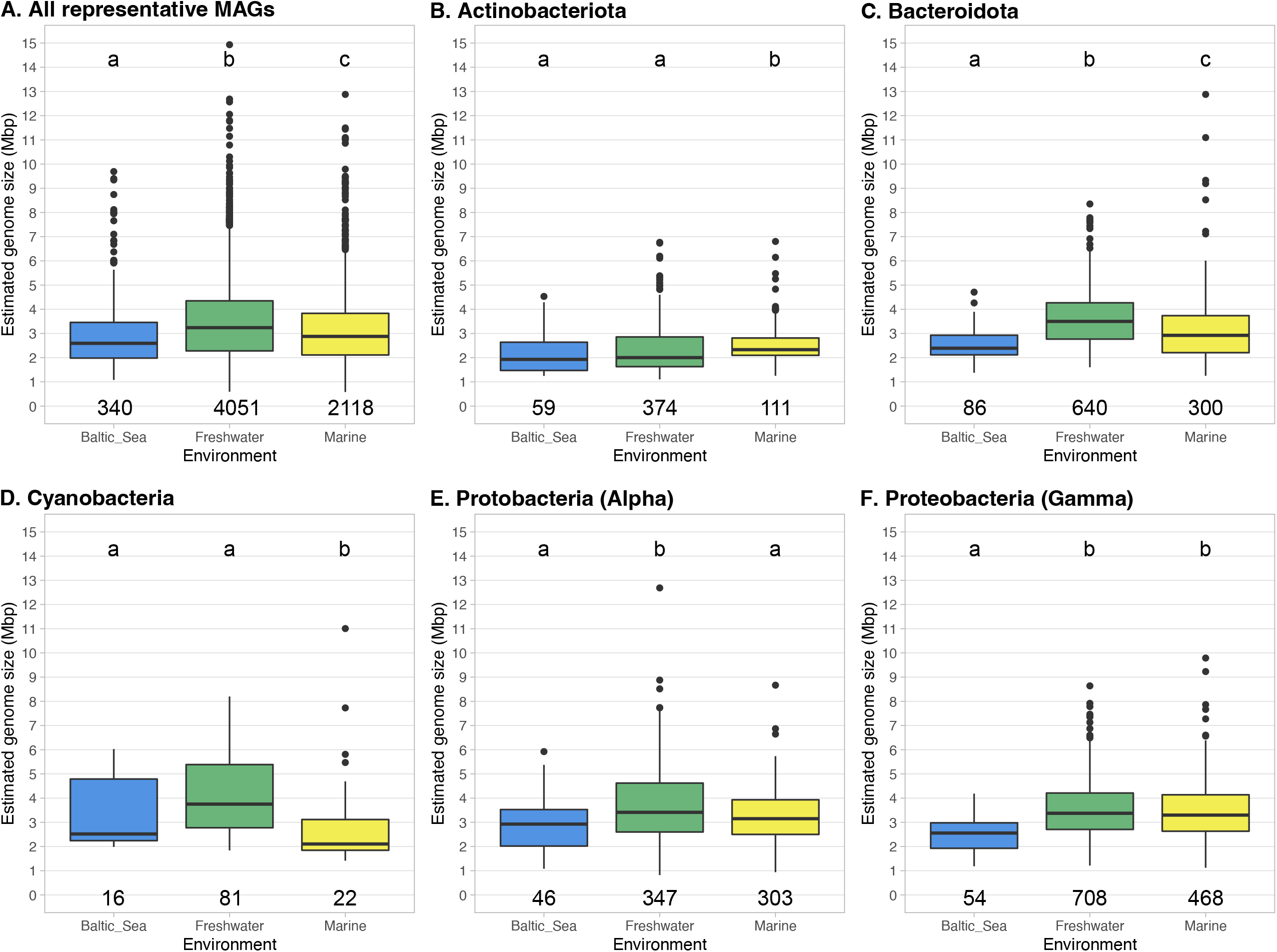
Overview of the estimated genome size of pelagic Bacteria and Archaea obtained from Baltic Sea (blue), freshwater (green) and marine (yellow) using only representative MAGs (calculates using 95% average nucleotide identity) with >75% completeness. We compare all representative MAGs (**Panel A**), only phylum Actinobacteriota (**Panel B**), phylum Bacteroidota (**Panel C**), phylum Cyanobacteria (**Panel D**), class Alphaproteobacteria (**Panel E**) and class Gammaproteobacteria (**Panel F**). Different letters indicate significant differences p < 0.05 (Kruskal-Wallis non-parametric test; multiple testing corrected with Benjamini-Hochberg). Numbers below the boxes indicate the number of MAGs per environment.

We then divided all MAGs into phyla and focus on the most common from aquatic environments: phyla Actinobacteriota, Bacteroidota, Cyanobacteria and Proteobacteria (classes Alpha and Gammaproteobacteria) (**Figure 4B-F**). We test if the genome size differences are consistent across phyla. We observe that only phylum Bacteroidota follows the same trend as the full dataset in the average estimated genome size (**Figure 4C**). On the other hand, MAGs from phylum Actinobacteriota present the biggest genome sizes for marine environments, while no difference on average genome size is observed between freshwater and brackish (**Figure 4B**). This result updates previous observations on aquatic Actinobacteriota genome size variation (23). Opposite to Actinobacteriota, MAGs from phylum Cyanobacteria show the smallest average genome size for marine environments, while we do not observe statistical differences between brackish and freshwater MAGs (**Figure 4D**). Similar trends were observed for isolates and MAGs of picocyanobacteria (22,49). However, it is important to remark that DNA extraction, assembly, binning and/or quality check of aquatic cyanobacterial MAGs is still a big challenge that needs to be addressed (**Supplementary Material 2**) (50,51). All in all, these results hint at the complicated ecological role of genome size in pelagic bacterial groups, where environment (2,15,16), lifestyle (4,13,14) and taxonomy (11,12) are intertwined.

### Smaller bacterial genomes in the Baltic Sea tend to lack certain functional categories

We selected the bacterial MAGs with >90% completeness for metabolic annotation and analyze which functional categories correlate with genome size in brackish sediments and water column (**Figure 5 A-R** and **Supplementary Material 3**). Functional categories include different but related metabolic pathways (KEGG modules), and each module comprises multiple specific metabolic reactions (module steps) (52). In our results, we observe patterns of negative correlation between estimated genome size and number of module steps per Mbp in most of the functional categories analyzed (**Figure 5**). For example, given the core functions amino acid metabolism, aminoacyl tRNAs and central carbohydrate metabolism, we observe a higher number of module steps per Mbp at lower genome sizes in both environments both in pelagic and benthic MAGs (**Figures 5A, D** and **J**). However, genome size does not explain the number of module steps per Mbp in all functional categories and metabolisms, especially in the case of non-essential functions such as drug resistance and transport systems (**Supplementary Material 3F** and **N**). These results suggest that streamlining of genomes select for specific functions and not the whole genome. This could be explained by the Black Queen Hypothesis (53); when the fitness cost of a function is higher than its benefit, microbes might lose it and, instead, obtain benefit from leaky metabolites from neighboring cells, establishing interdependencies. The loss of those specific functions in the Baltic Sea might have consequential long-lasting metabolic partnerships within the community.

**Figure 5.**
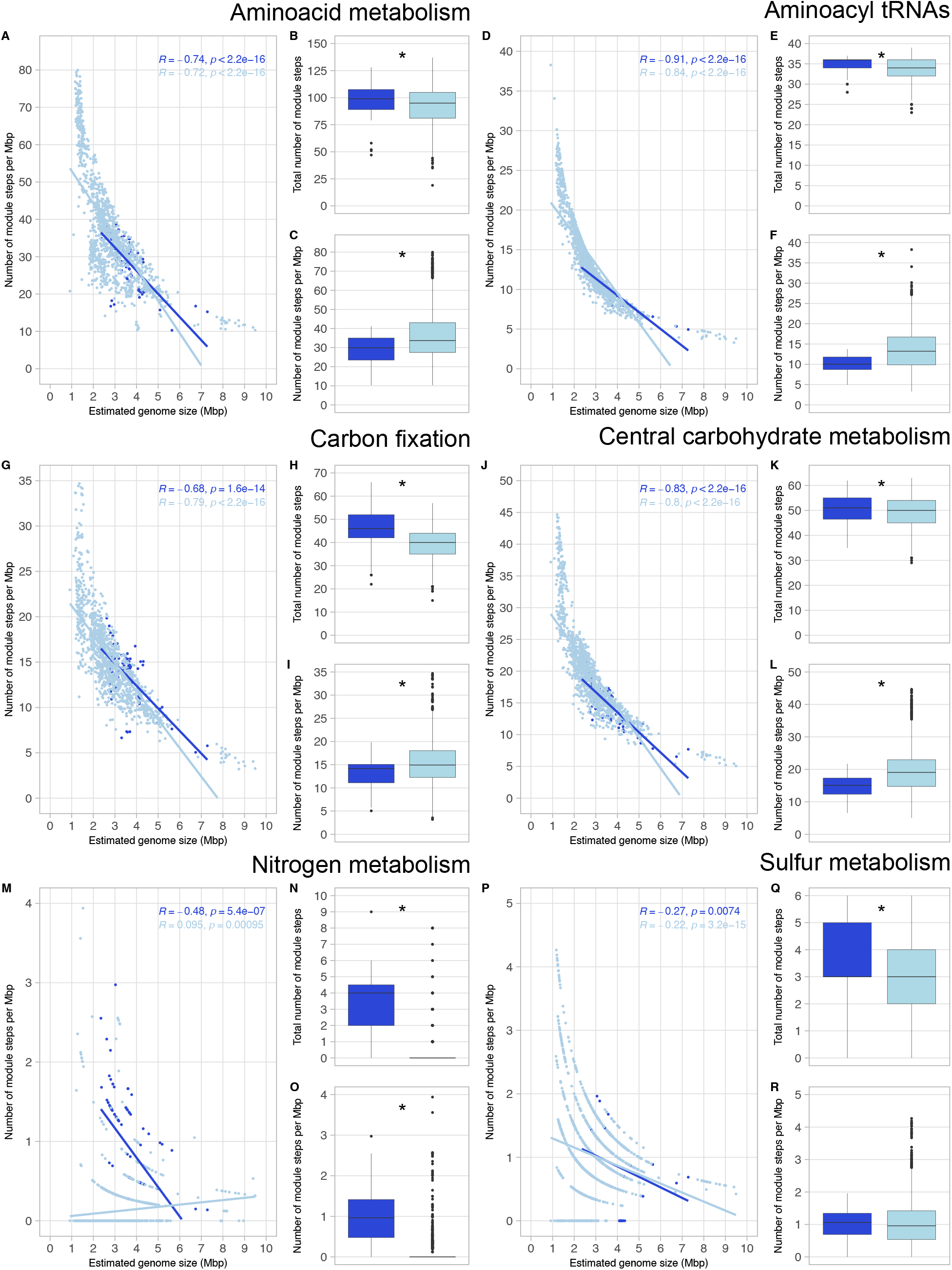
Overview of the presence of module steps for six metabolic categories: amino acid metabolism (**Panels A-C**), aminoacyl tRNAs (**Panels D-F**), carbon fixation (**Panels G-I**), central carbohydrate metabolism (**Panels J-L**), nitrogen metabolism (**Panels M-O**) and sulfur metabolism (**Panels P-R**). We used all bacterial MAGs with very-high quality (>90% completeness and <5% contamination), from both environments (99 MAGs from sediments and 1215 MAGs from water column). Stars in boxplots indicate significant differences p < 0.05 (Wilcoxon non-parametric test). **Panels B, E, H, K, N** and **Q** indicate total number of module steps. **Panels C, F, I, L, O** and **R** indicate the number of module steps per Mbp.

When considering the total number of module steps, we observe that genomes of benthic bacteria code for a higher number of module steps than pelagic bacteria in amino acid metabolism, aminoacyl tRNAs, carbon fixation, nitrogen and sulfur metabolism (**Figures 5B, E, H, N** and **Q**) (Wilcoxon test, *p* < 0.01). This could be the result of a confounding effect of genome size: microorganisms from sediments have bigger genome sizes than those in the water column (**Figure 2**) and therefore, code for a higher number of functions. Hence, the total number of module steps per Mbp in all six categories was analyzed. If streamlining affects all functional categories similarly, coding density trends would be similar for all functional categories. Indeed, we observe that pelagic bacteria present a greater number of module steps per Mbp than sediment bacteria in amino acid metabolism (**Figure 5C**), aminoacyl tRNAs (**Figure 5F**), carbon fixation (**Figure 5I**) and central carbohydrate metabolism (**Figure 5L**). However, nitrogen metabolism shows the opposite trend: sediment bacteria have a greater number of module steps per Mbp than water column bacteria (**Figure 5O)**. Literature report that Baltic sediments contain about 95% of the total pool of nitrogen while the water column only 5%, hence the water column only carries a small part of the overall nitrogen cycle (54). As mentioned above for carbon, a higher availability of resources, including nitrogen-derived compounds, would also imply a lower evolutionary pressure for streamlined genome sizes in Baltic sediments. These results indicate that benthic bacteria potentially play a bigger role than pelagic bacteria in nitrogen cycling of autotropic systems like the Baltic Sea (55,56).

Although most of the MAGs presented at least 1 module step related to drug resistance (96.97% sediment MAGs and 98.81% pelagic MAGs), bacteria from sediments presented a higher number of antibiotic resistance module steps than pelagic bacteria. This applies both to total number of module steps and the number of module steps per Mbp (**Supplementary Material 3**). Our results confirm that aquatic sediments are reservoirs of antibiotic resistance genes (57,58). Just as sediments harbor more than double the organic carbon than the water column, this allows microbial genomes to have a bigger size and code for a higher number of genes. This allows microbes to upkeep non-essential functions and allow metabolic reservoir in the sediments.

### Baltic sediments and water column harbor bacteria with different metabolic capabilities

From all >90% completeness MAGs, we selected only bacterial phyla with five or more high-quality MAGs to observe how metabolism differs between different taxa in Baltic sediments and water column (**Figure 6**).

**Figure 6.**
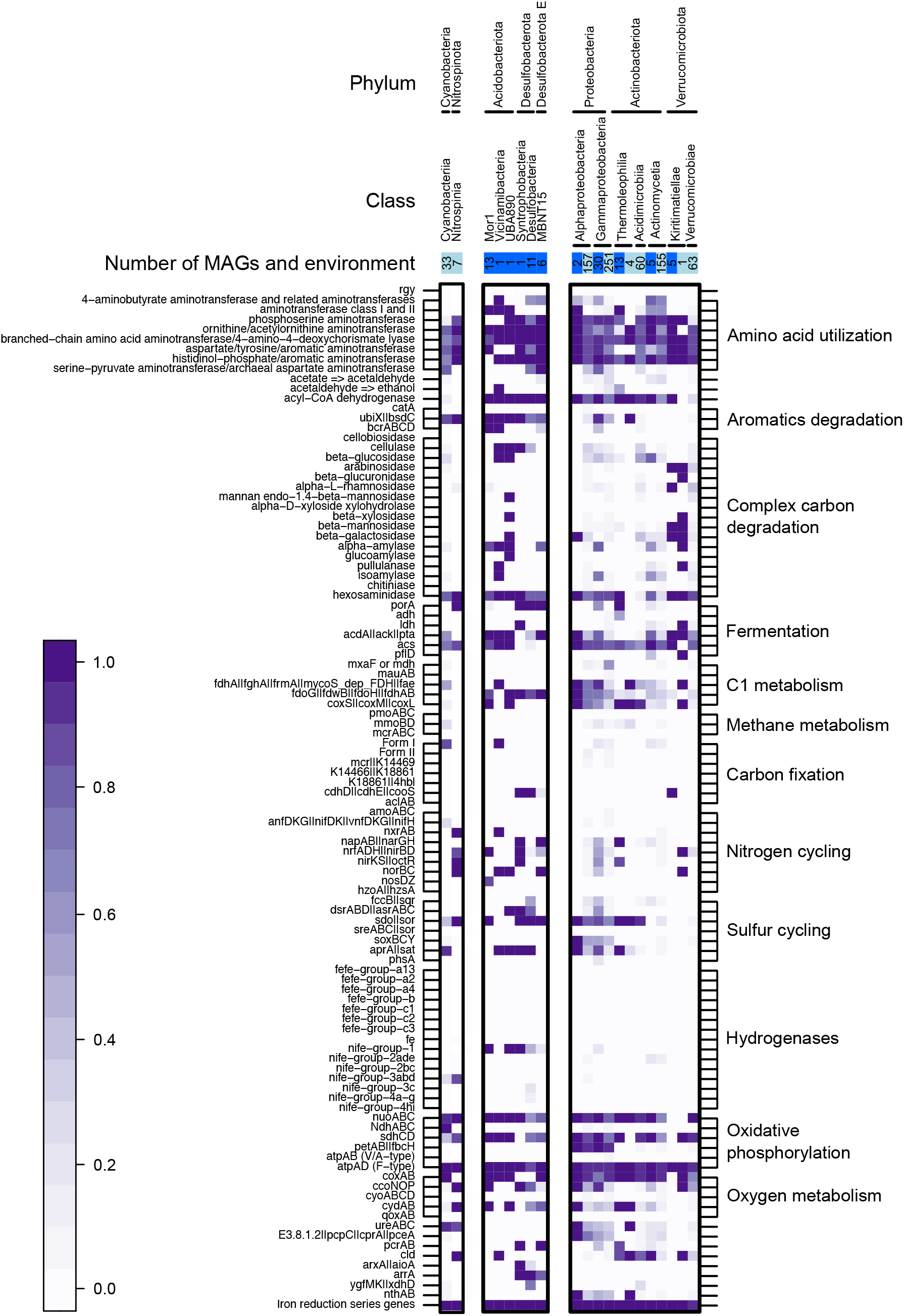
Metabolic potential of all high-quality MAGs (>90% completeness and <5% contamination). We selected all phyla with at least 5 MAGs, and then divided by class. Boxes on the top of the figure indicate environment (pelagic in light blue and sediments in dark blue), and inner numbers indicate the number of MAGs per category. In the heatmap, white squares indicate absence of a given gene in all MAGs, and the darkest purple indicates presence in all of them (gradient scale on the bottom-left part of the figure for reference).

We observed the presence of the genes *cdhH* | *cdhE* | *cooS* from the Wood-Ljungdhal pathway exclusively in marine sediments (**Figure 6**), particularly in phyla Desulfobacterota, Desulfobacterota E and Verrucomicrobiota (class Kiritimatiellae). This is a carbon fixation pathway predominant in acetogenic bacteria found in anoxic conditions (59). Complementarily, we also find putative fermentation genes for acetogenesis (*acdA* | *ack* | *pta*) to be widespread across taxa in Baltic sediments. This would explain the potential success of acetogenic metabolism in brackish sediments (60). We also find that the *acs* gene for acetate fermentation into acetyl-CoA is widely distributed in both sediments and water column. These results support the common distribution of acetogens and the Wood-Ljungdhal pathway in Baltic glacial sediments (61).

No FeFe hydrogenases were detected, but different NiFe hydrogenases were spotted to differ between environments: NiFe groups 3abd were detected mainly in pelagic bacteria, while NiFe group 1 in sediments (**Figure 6**). NiFe group 1 hydrogenases could be playing a vital role in nitrate (NO3-), sulfate (SO42-) and iron (Fe^3+^) reduction: these molecules can act as acceptors of electrons coupled to H2 oxidation in anoxic conditions (62). Moreover, putative genes coding for the reduction of the above-mentioned molecules were also detected on our sediment dataset (*napAB* | *narGH* for nitrate reduction, *aprA* | *sat* for sulfate reduction, and iron reduction series genes). These results suggest a key role of sediment bacteria in sulfur, nitrogen, and iron cycling in the Baltic Sea. For example, sulphate reducers such as Desulfobacterota found in our benthic MAGs collection, most likely contribute to the release of Fe-bound phosphorus from sediments to the water column (63).

### Conclusions

In this research article, we provide a comprehensive analysis to investigate the ecological implications of microbial genome in the Baltic Sea. We show that genome size in Bacteria and Archaea is linked to the environment (**Figures 2, 3, 4** and **S1** and **Supplementary Material 1**), taxonomic identity (**Table 1** and **Figure 3**) and metabolic potential (**Figure 5** and **Supplementary Material 3**). We also provide some insights on how distinct pelagic and benthic microbial communities in the Baltic Sea are: not only microbial MAGs retrieved from these two environments are different in taxonomy (**Table 1**), but also in genome size (**Figure 3**) and metabolism (**Figures 5** and **6**). This highlights water bodies are highly heterogeneous biomes, with highly distinct microbial communities between micro-niches. With the continuous progression of aquatic microbial ecology and the development of new isolation, omics and bioinformatic techniques, future research should provide a more complete and unbiased view of genome sizes distribution in nature and its ecological implication in microbial populations.

## MATERIAL AND METHODS

### Baltic Sea metagenomes collection

For this study we compiled new and public Baltic Sea metagenomes from the water column and the sediments. The final dataset consisted of 219 metagenome samples from a wide range of locations in the Baltic Sea that include 5 independent datasets (**Table S1** and **Figure 1**). We compiled 108 sediment metagenomes, of which 59 were collected in 2019 and recently published (29), and we collected 49 metagenomes from 2016 to 2018. The pelagic dataset consists of 118 pelagic samples collected from 2011 to 2015 published in three different studies (46,64,65). All five datasets have abiotic metadata of depth (m), salinity (PSU), temperature (C) and oxygen concentration (mg/L) (**Table S1**).

### Environmental sampling

The top 2 cm of sediment was collected at soft bottom clay-muddy habitats from 59 stations from north to south in the Baltic Sea in 2019, following the sampling described in Broman et al., (2022) (**Table S1** for coordinates). Briefly, one sediment core was collected per station using a Kajak gravity corer (surface area: 50 cm^2^, one core per station) and the top 0–2 cm layer was sliced into a 215 ml polypropylene container (207.0215PP, Noax Lab, Sweden). The sediment was homogenized and stored at -20°C on the boat, kept on an iced cooler without thawing for ∼2 h during transportation to the university, and finally stored again at -20 °C until DNA extraction. Bottom water (∼20 cm above the sediment surface) was collected at each station with a Niskin bottle. This was followed by on deck measurements of bottom water temperature, salinity, and dissolved O2 using a portable multimeter (HQ40D, Hach).

### DNA extraction and sequencing

The sediment samples were thawed, homogenized, and a subsample of 0.25 g was used for DNA extraction using the DNeasy PowerSoil kit (Qiagen) according to the manufacturer’s protocol. The quantity and quality of eluted DNA were measured using NanoDrop One spectrophotometer and Qubit 2 (both by ThermoFisher Scientific) to ensure that samples meet the minimum requirements for sequencing. The samples were then sequenced at the Science for Life Laboratories facility on one NovaSeq 6000 S4 lanes using a 2 × 150 bp setup. Sequencing yielded on average 53.0 million reads per sample.

### MAGs collection

Assembling and binning of the 108 sediment metagenomes resulted in 2248 bins. To obtain bins from metagenomes, we followed the 0053_metasssnake2 pipeline (https://github.com/moritzbuck/0053_metasssnake2) (v0.0.2). In this pipeline we used Sourmash (66) to create signatures, Megahit (67) to obtain single-sample assemblies and Metabat2 (68) for the binning of the assemblies. We used default parameters throughout the whole pipeline. Quality of the bins was assessed using CheckM (v1.1.3) (69): we used the typical workflow (*lineage_wf*) with default parameters, and selected only bins with quality of completeness >75% and contamination <5%. From all the bins, only 216 passed our quality threshold and we named those MAGs (metagenome-assembled genomes). All MAGs were taxonomically classified using GTDB-tk (v1.5.0) (70) according to the GTDB classification (data version r202) (71). The quality of the MAGs belonging to phyla Actinobacteriota and Patescibacteria were assessed separately using a custom set of marker genes. Preliminary quality check of Actinobacteriota genomes in a publicly available freshwater dataset (47) show that default parameters underestimate the quality of the MAGs that are classified as Actinobacteriota compared to using a custom marker gene set (**Supplementary Material 2**). We obtained the custom marker gene set using Ac1 circular genomes (72,73). In the case of phylum Patescibacteria, we used a custom set of maker genes provided by CheckM (69,74).

Complementarily, we used 1920 pelagic MAGs that were published (46) and passed the >75% completeness and <5% contamination threshold. All high-quality MAGs from the sediments and water column were dereplicated using fastANI (95% ANI threshold as estimator of genetic unit) and mOTUlizer (v0.3.2) (75,76). From the 216 with >75% completeness MAGs, 56 were chosen as representatives. From the 1920 pelagic MAGs, 340 were chosen as representatives. All genomic information for pelagic and benthic MAGs is included in **Table S2**.

### Genome size analysis

We studied genome size in two different levels: entire microbial community and bacterial/archaeal MAGs. To study differences in genome size between Baltic sediments and water column at the community level, we first calculated the average genome size (AGS) of the metagenomes using MicrobeCensus (v1.1.0) (77). MicrobeCensus estimates the AGS of a microbial community from metagenomes by aligning reads to a set of single-copy genes that are widely distributed across taxa to calculate their abundance, with highly accurate estimations (41). We excluded one of the pelagic datasets (65) due to the presence of spike-in DNA. We used default parameters, but we set the number of reads sampled to 10 million (*-n 10 000 000*). To estimate the genome size of the microbial MAGs, we divided the MAGs assembly size by the completeness (provided by CheckM, ranging from 0 to 1). To study genome size variation between our pelagic brackish dataset and other major aquatic environments (freshwater and marine), we compiled the metagenomic information from all pelagic MAGs from marine and freshwater environments (>75% completeness and <5% contamination) from (2) (**Table S3**).

### Metabolic annotation

To analyze the metabolic potential of sedimentary and pelagic bacteria, we selected all MAGs with completeness >90% and contamination <5%. In total, we obtained 99 MAGs from sediments and 1241 MAGs from the water column. The metabolic potential of sediment and pelagic MAGs was reconstructed using Prodigal annotations (v2.6.3) (78). We used the resulting protein translation files to predict biogeochemical and metabolic functional traits using METABOLIC (v4.0) (52). We used the METABOLIC-G script, using default settings.

### Statistical analysis

We performed Wilcoxon non-parametric test to analyze if there were significant differences between pairs of boxplots (**Figures 2, 3** and **5** and **Supplementary Material 3**). Asterisks in boxplots indicate significant differences p < 0.05. In **Figures 3B** and **4** we performed Kruskal-Wallis test corrected with Benjamini-Hochberg, to test statistical differences between groups. Different letters are the result of this non-parametric test; p < 0.05. We performed a ANOVA type II analysis to test the effect of abiotic factors and their interactions on the AGS, using the function *aov* from the R package *stats* v3.6.3 (79) (**Supplementary Material 1**). We obtained the correlation coefficients on scatterplots (**Figures 5** and **S1**) to test the fit of our data to linear regressions using the function *stat_cor* from the R package *ggpubr* v0.4.0 (80).

## Data availability

All metagenome datasets are available in public repositories under NCBI project accession number SRP077551 and ENA accession numbers PRJEB34883, PRJEB22997 and PRJEB41834. Specific accession numbers of all metagenomes are available in **Table S1**. Pelagic MAGs can be found on project PRJEB34883 and benthic MAGs on project PRJNA891615. Assembly and binning of the dataset provided in this paper used scripts available at https://github.com/moritzbuck/0053_metasssnake2. Supplemental material can also be accessed 10.17044/scilifelab.21378294.

## ACKNOWLEDGEMENTS

We thank Elias Broman for his help on the acquisition of the benthic dataset of metagenomes and metadata and Ola Svensson, Caroline Raymond and Jonas Gunnarsson for assistance during sampling. We also thank Juanita Gutiérrez-Valencia for her suggestions in statistical analysis.

This work was supported by SciLifeLab. The authors acknowledge support from the National Genomics Infrastructure in Stockholm funded by Science for Life Laboratory, the Knut and Alice Wallenberg Foundation and the Swedish Research Council, and SNIC/Uppsala Multidisciplinary Center for Advanced Computational Science for assistance with massively parallel sequencing and access to the UPPMAX computational infrastructure. Computational work and data handling were enabled by resources in the projects SNIC 2020/5-159, 2021/5-133, 2022/5-137, 2020-6-60 and 2022/6-77 provided by the Swedish National Infrastructure for Computing (SNIC) at UPPMAX, partially funded by the Swedish Research Council through grant agreement no. 2018-05973.

AR-G and SLG conceptualized and designed research project. AR-G, MB and SLG refined the project idea. AR-G, AFA and FJAN participated in data collection. AR-G and DIS did the molecular work. AR-G and MB performed bioinformatic analysis to obtain MAGs from sediment raw sequences. AR-G and SLG performed data analysis. AR-G and SLG drafted the first manuscript. AR-G, DIS and SLG did literature searches. All authors contributed to the writing and editing of the manuscript.

## CONFLICTS OF INTEREST

The authors declare that there are no competing interests.

## Notes

### Competing Interest Statement

The authors have declared no competing interest.

### Summary of Updates

Abstract, introduction, results and discussion and conclusions updated to clarify. Figure S2 included. Order of supplemental files updated.

