## Supplementary figures and images for "Linking prokaryotic genome size variation to metabolic potential and environment"

### Figure S1

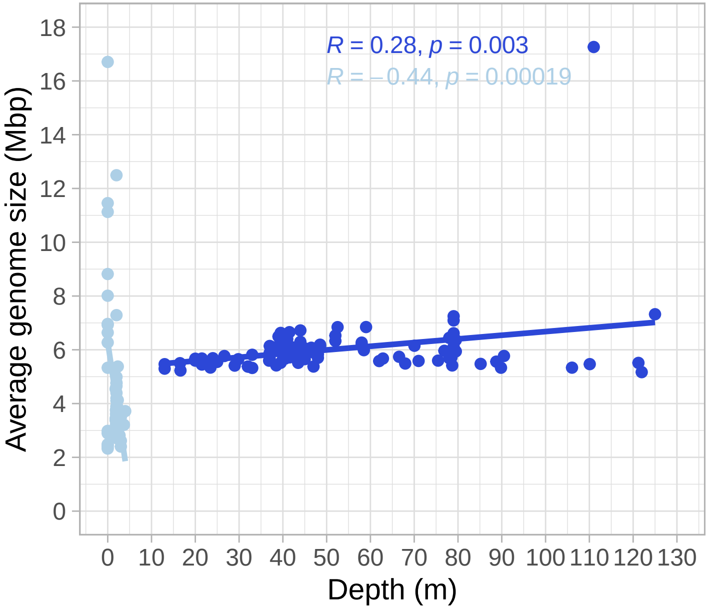

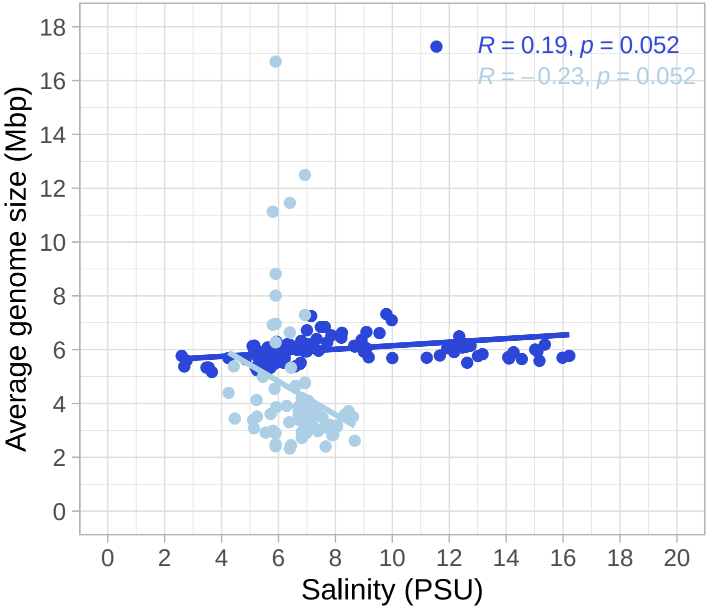

### Figure S2

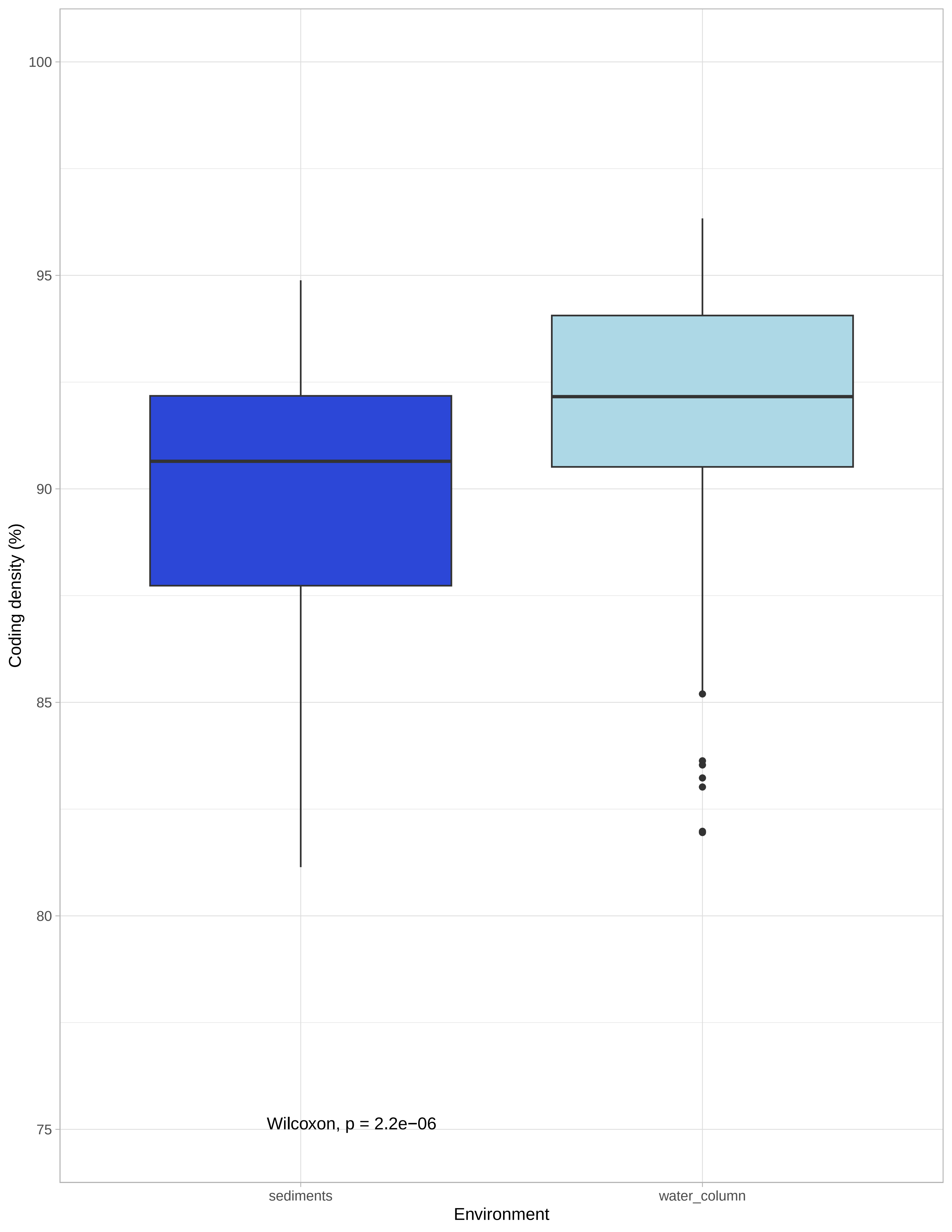

### Supplementary Material 1

**Supplementary material 1**

Water column

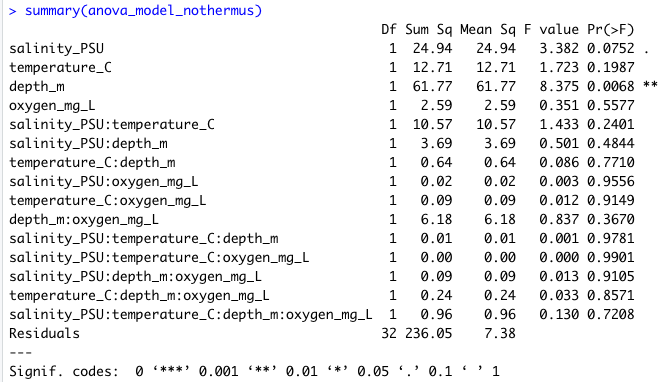

Sediments

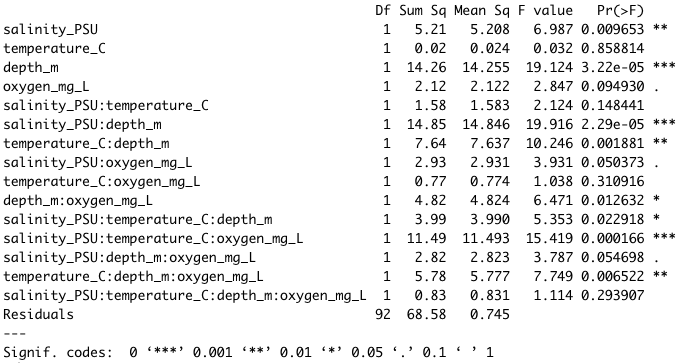
