## Supplementary Material 2 for "Linking prokaryotic genome size variation to metabolic potential and environment"

**Supplementary Material 3**

Table: Full freshwater actinobacterial genomes used to calculate the lineage markers in CheckM.

| **GenBank Accession** | **Lake** | **Tribe** | **Assembly size (Mb)** | **GC content (%)** | **Citation** |
| --- | --- | --- | --- | --- | --- |
| CP015603 | Soyang | acI-A1 | 1.35 | 49.05 | Kang et al. 2017 |
| CP016770 | Zurich | acI-A1 | 1.36 | 47.92 | Neuenschwander et al. 2018 |
| CP016772 | Zurich | acI-A1 | 1.37 | 47.96 | Neuenschwander et al. 2018 |
| CP016773 | Zurich | acI-A1 | 1.34 | 48.56 | Neuenschwander et al. 2018 |
| CP016774 | Zurich | acI-A1 | 1.33 | 48.26 | Neuenschwander et al. 2018 |
| CP016775 | Zurich | acI-A1 | 1.33 | 48.18 | Neuenschwander et al. 2018 |
| CP016777 | Zurich | acI-A1 | 1.35 | 47.98 | Neuenschwander et al. 2018 |
| CP016778 | Zurich | acI-A1 | 1.33 | 48.22 | Neuenschwander et al. 2018 |
| CP016781 | Zurich | acI-A1 | 1.27 | 48.31 | Neuenschwander et al. 2018 |
| CP015604 | Soyang | acI-A4 | 1.46 | 46.96 | Kang et al. 2017 |
| CP016769 | Zurich | acI-A4 | 1.47 | 47.54 | Neuenschwander et al. 2018 |
| CP016780 | Zurich | acI-A4 | 1.39 | 47.78 | Neuenschwander et al. 2018 |
| CP016783 | Zurich | acI-A4 | 1.42 | 47.76 | Neuenschwander et al. 2018 |
| CP015605 | Soyang | acI-A7 | 1.51 | 45.45 | Kang et al. 2017 |
| CP016776 | Zurich | acI-A7 | 1.36 | 45.75 | Neuenschwander et al. 2018 |
| CP016782 | Zurich | acI-Phila | 1.33 | 45.02 | Neuenschwander et al. 2018 |
| CP016768 | Zurich | acI-B1 | 1.24 | 41.45 | Neuenschwander et al. 2018 |
| CP016771 | Zurich | acI-B1 | 1.22 | 42.37 | Neuenschwander et al. 2018 |
| CP016779 | Zurich | acI-B1 | 1.16 | 40.22 | Neuenschwander et al. 2018 |
| CP015606 | Soyang | acI-C1 | 1.55 | 51.31 | Kang et al. 2017 |

Isolate genomes in the table were used to recalculate the completeness of Actinobacteriota MAGs.

In the figure below we included all bins from stratfreshDB (Buck et al., 2021).

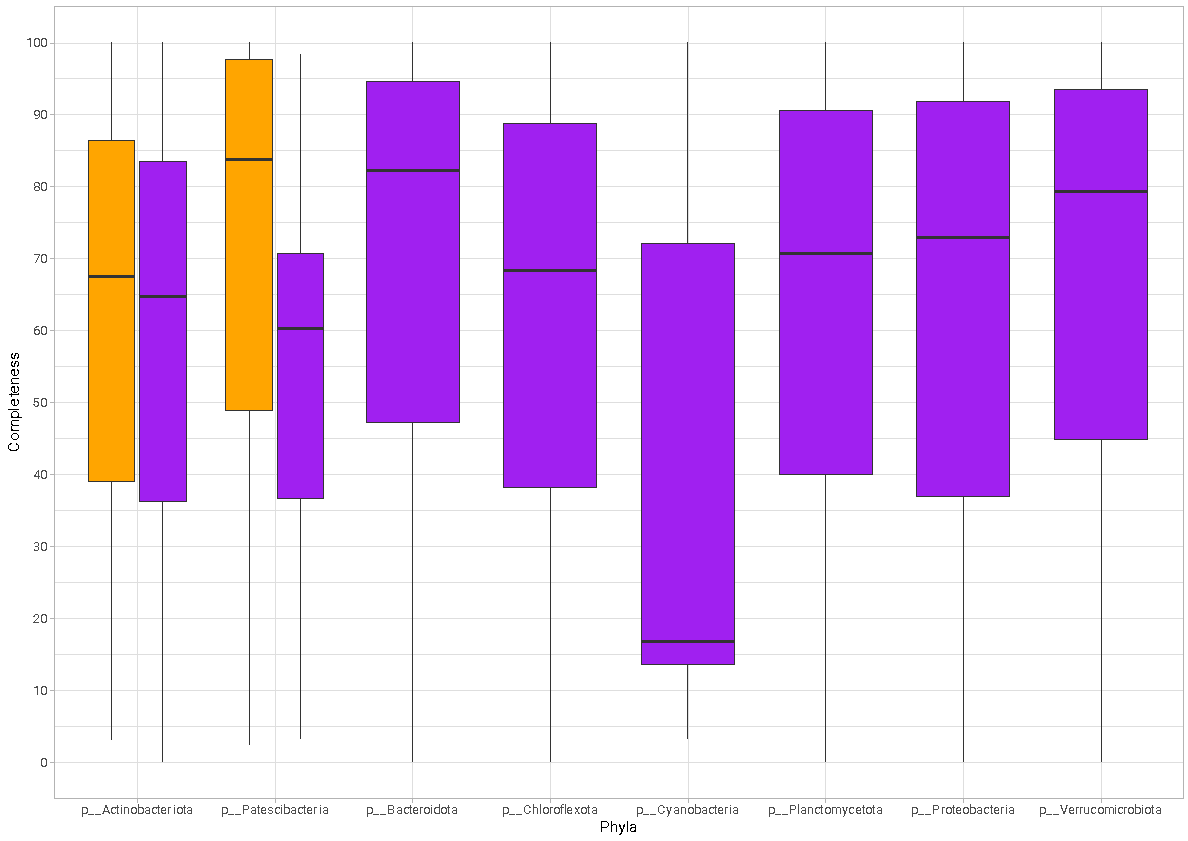

*	*

Completeness calculated with CheckM using default markers in purple

Completeness calculated with CheckM using specific markers for Actinobacteriota and Patescibacteria in orange.

* Denotes significant difference using Wilcoxon test

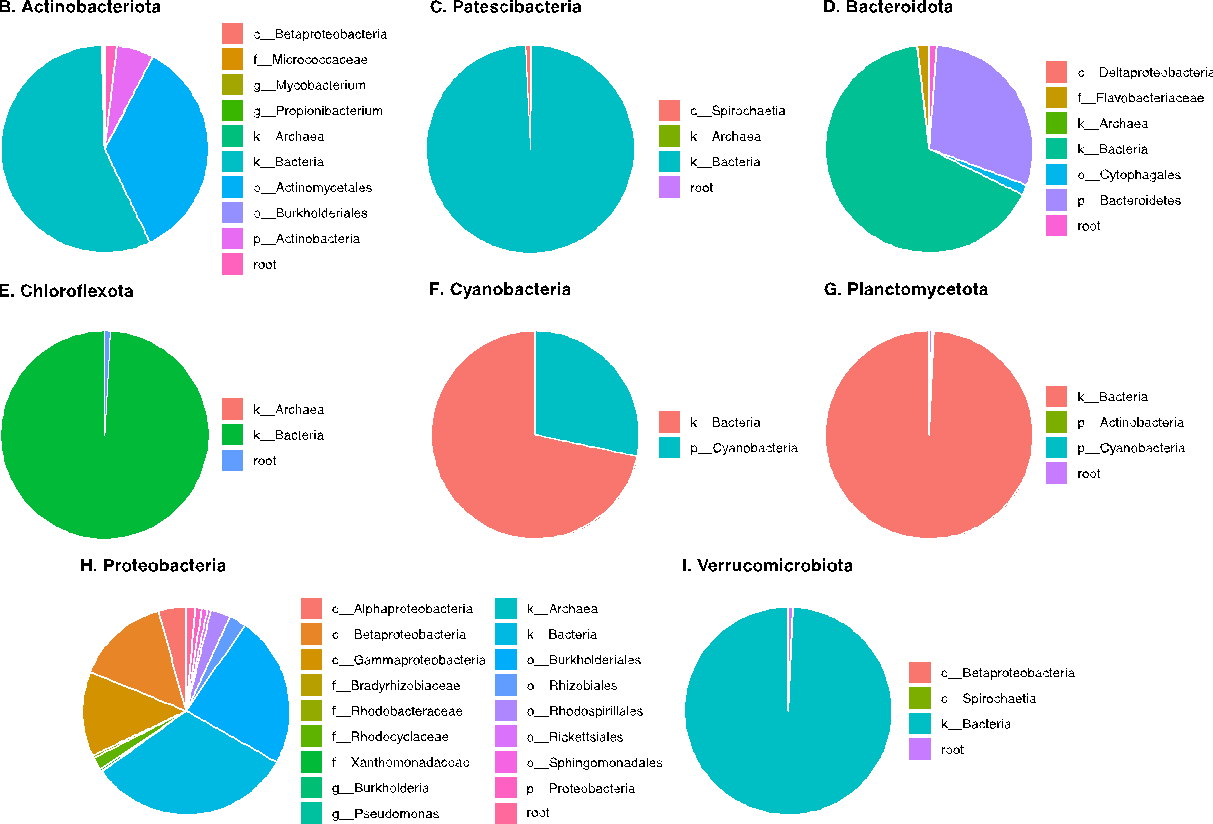

Lineage CheckM used to calculate completeness by default parameters.
