## Supplementary Material 3 for "Linking prokaryotic genome size variation to metabolic potential and environment"

**SUPPLEMENTARY MATERIAL 2**

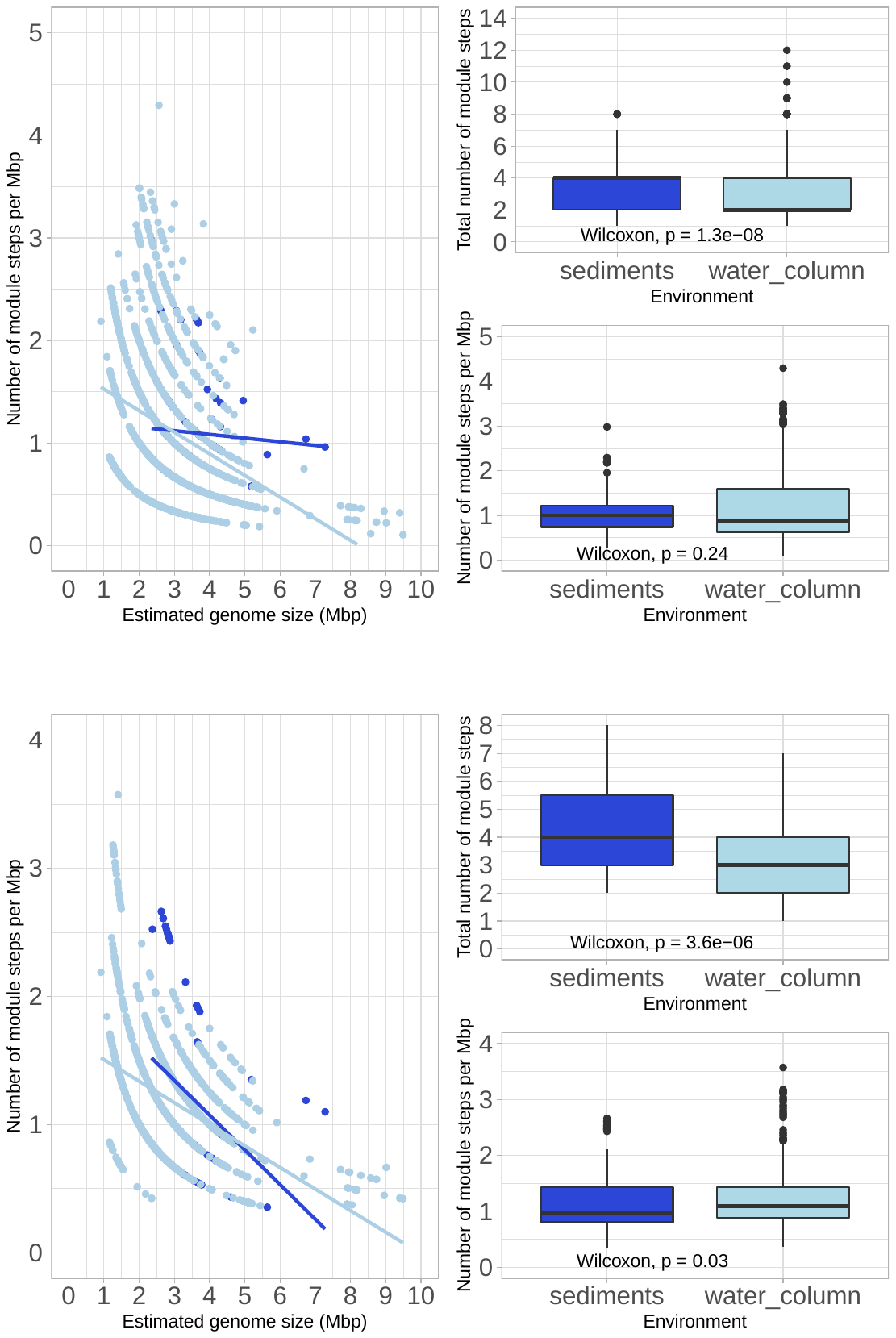

**Aromatics degradation**

y = 1.2 – 0.04x; R^2^ = 0.0033

y = 1.7 -0.21x; R^2^ = 0.12

*****

**Beta-lactam and macrolide biosynthesis**

y = 2.2 – 0.21x; R^2^ = 0.13

y = 1.7 – 0.17x; R^2^ = 0.19

*****

*****

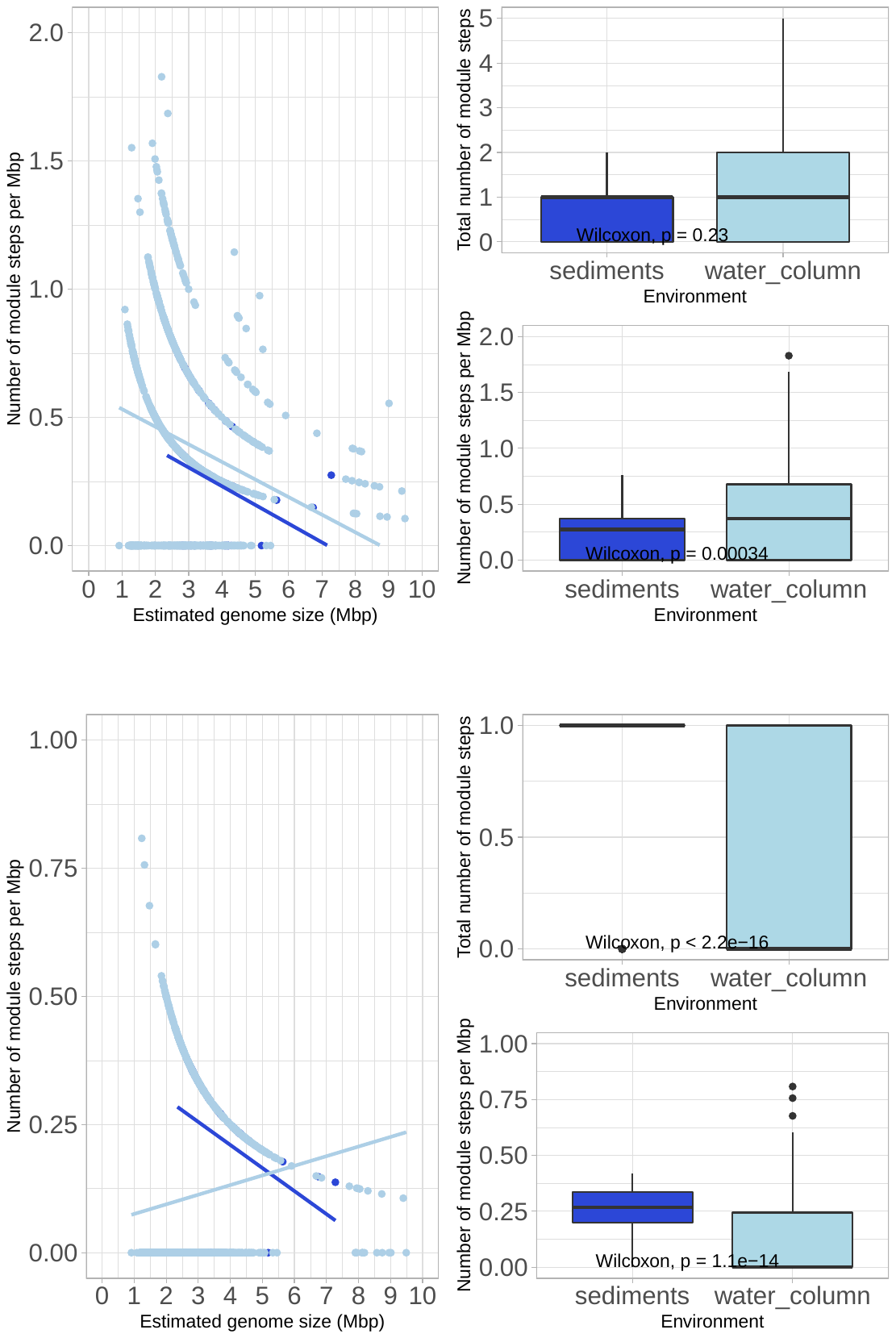
**Biosynthesis of secondary metabolites**

y = 0.52 – 0.073x; R^2^ = 0.072

y = 0.6 – 0.069x; R^2^ = 0.049

*****

**Cell signaling**

y = 0.39 – 0.045x; R^2^ = 0.08

y = 0.057 – 0.019x; R^2^ = 0.019

*****

*****

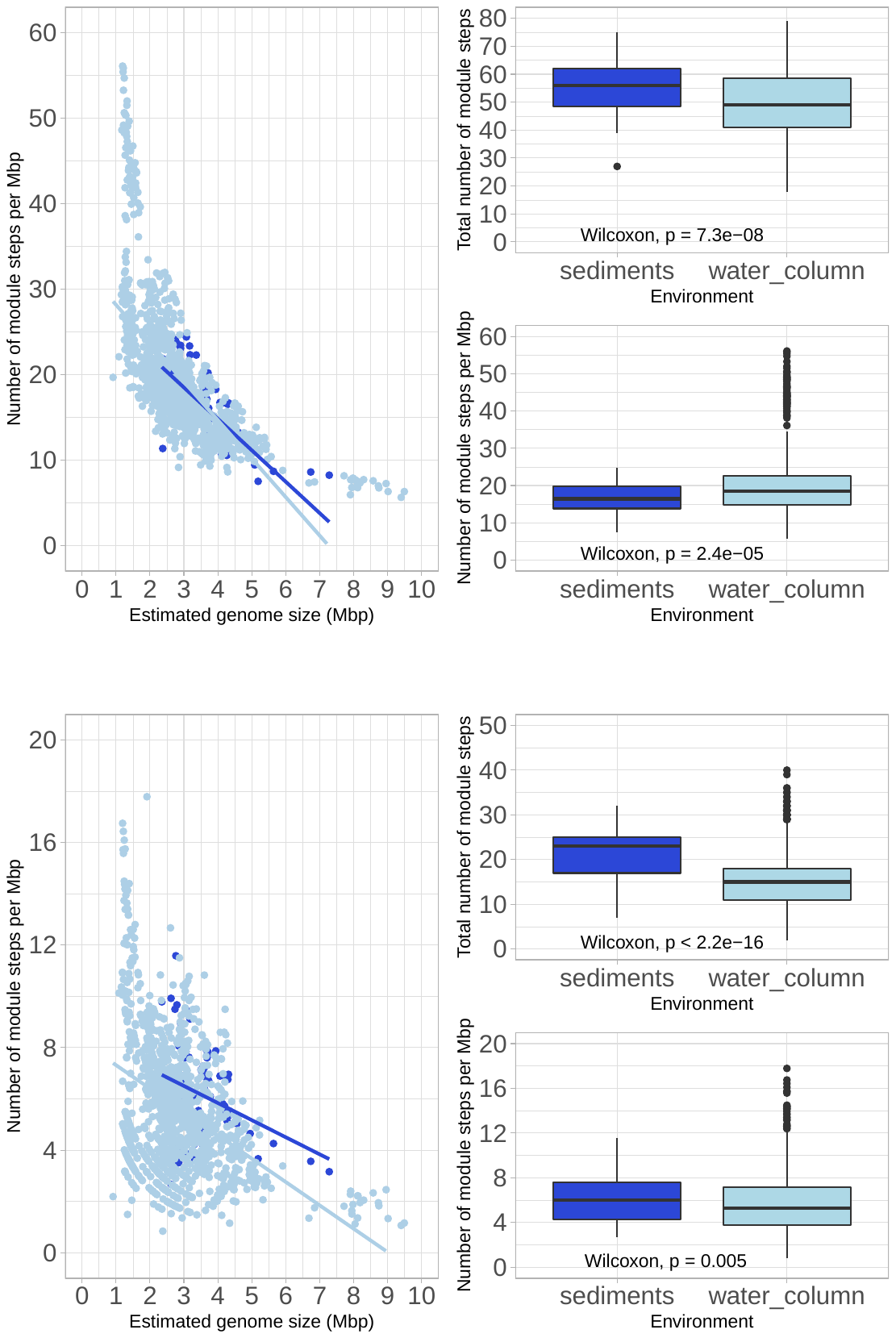
**Cofactor and vitamin metabolism**

y = 30 – 3.7x; R^2^ = 0.53

y = 33 – 4.5x; R^2^ = 0.45

*****

*****

**Drug resistance**

y = 8.5 – 0.67x; R^2^ = 0.083

y = 8.2 – 0.91x; R^2^ = 0.18

*****

*****

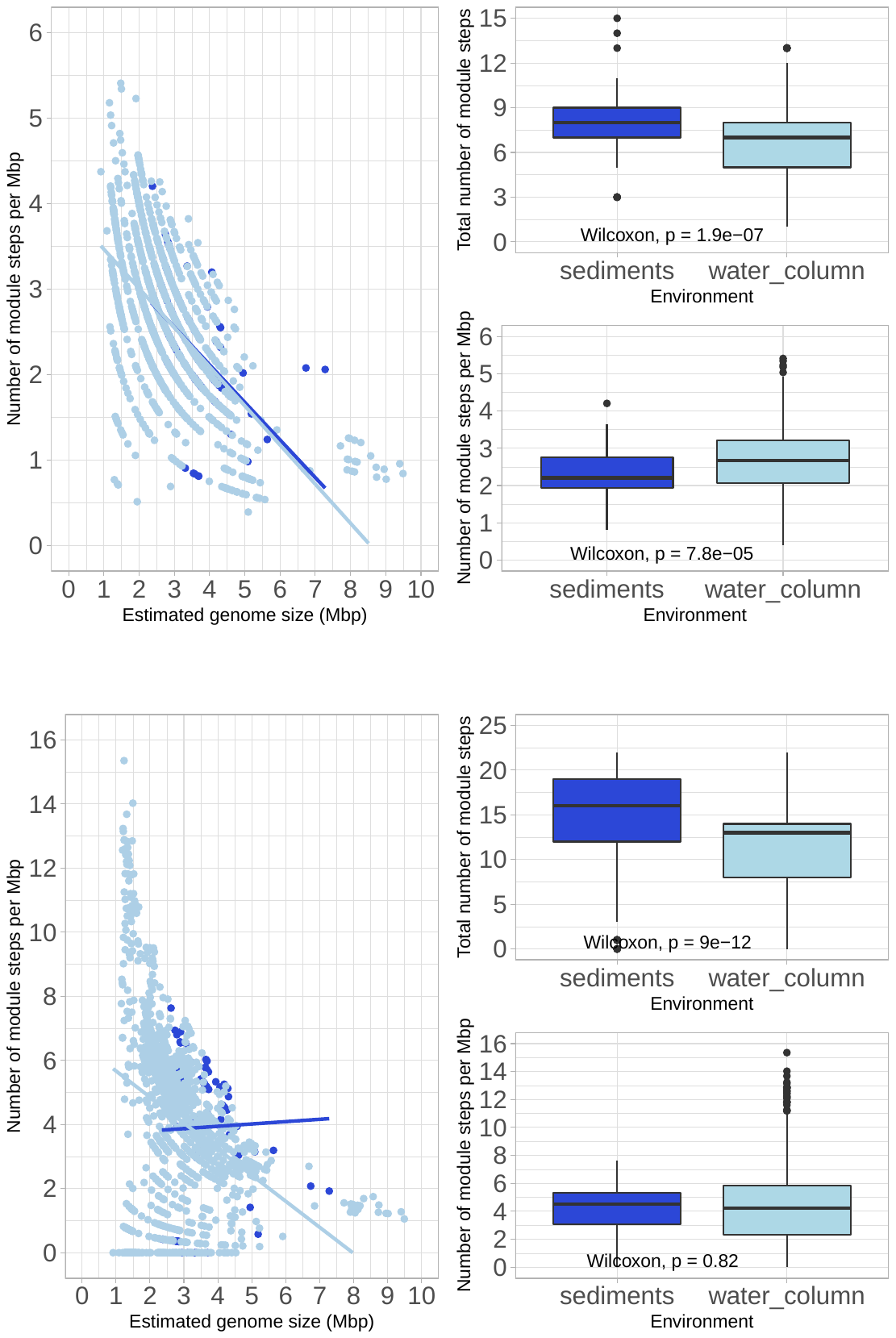
**Lipid metabolism**

*****

y = 3.9 – 0.073x; R^2^ = 0.29

y = 3.9 – 0.46x; R^2^ = 0.38

*****

**Lipopolysaccharides biosynthesis**

y = 3.7 + 0.073x; R^2^ = 0.001

y = 6.5 – 0.81x; R^2^ = 0.12

*****

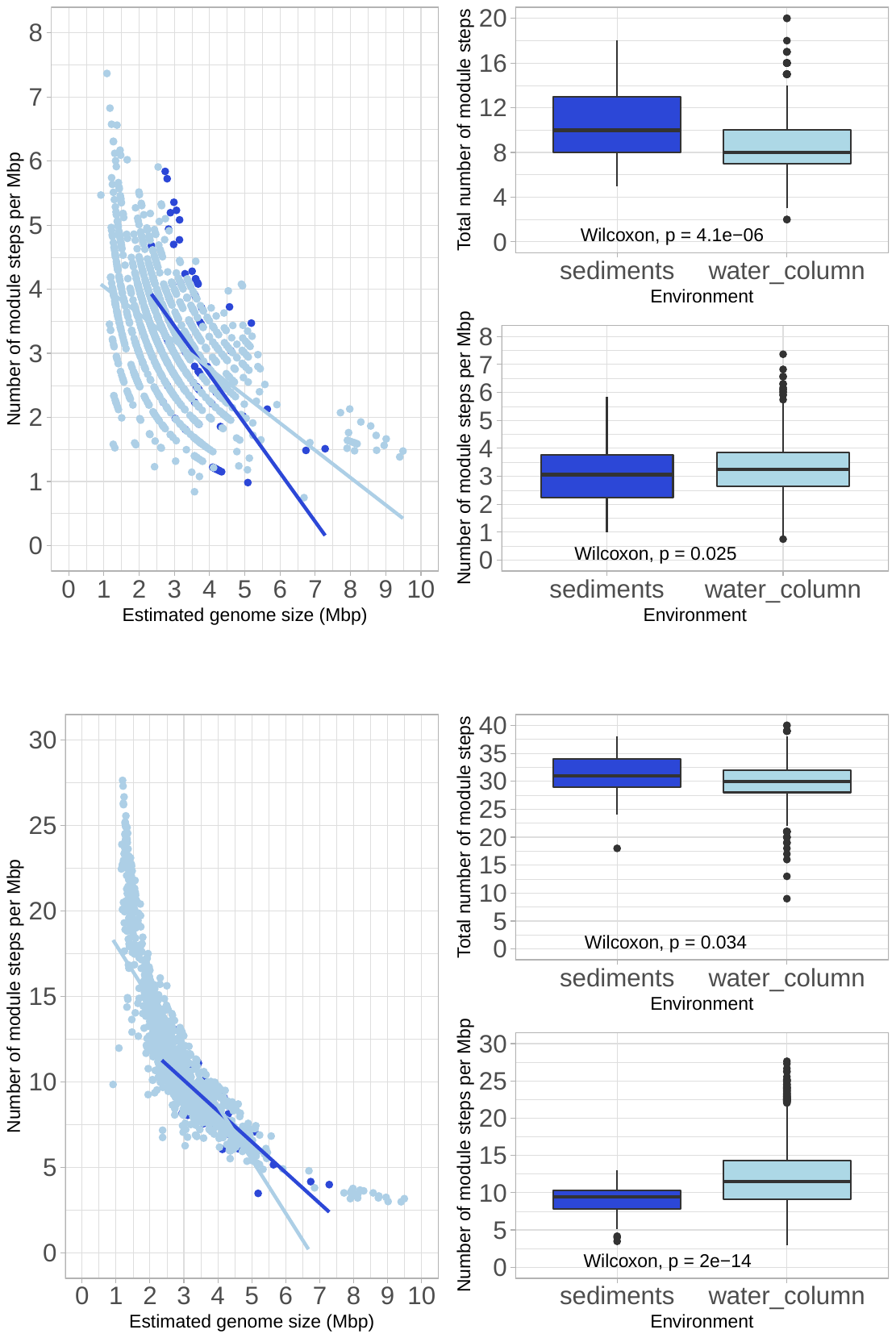
**Methane metabolism**

y = 5.7 – 0.076x; R^2^ = 0.3

y = 4.5 – 0.43x; R^2^ = 0.29

*****

*****

*****

*****

**Nucleotide metabolism**

y = 15 – 1.8x; R^2^ = 0.72

y = 21 – 3.1x; R^2^ = 0.67

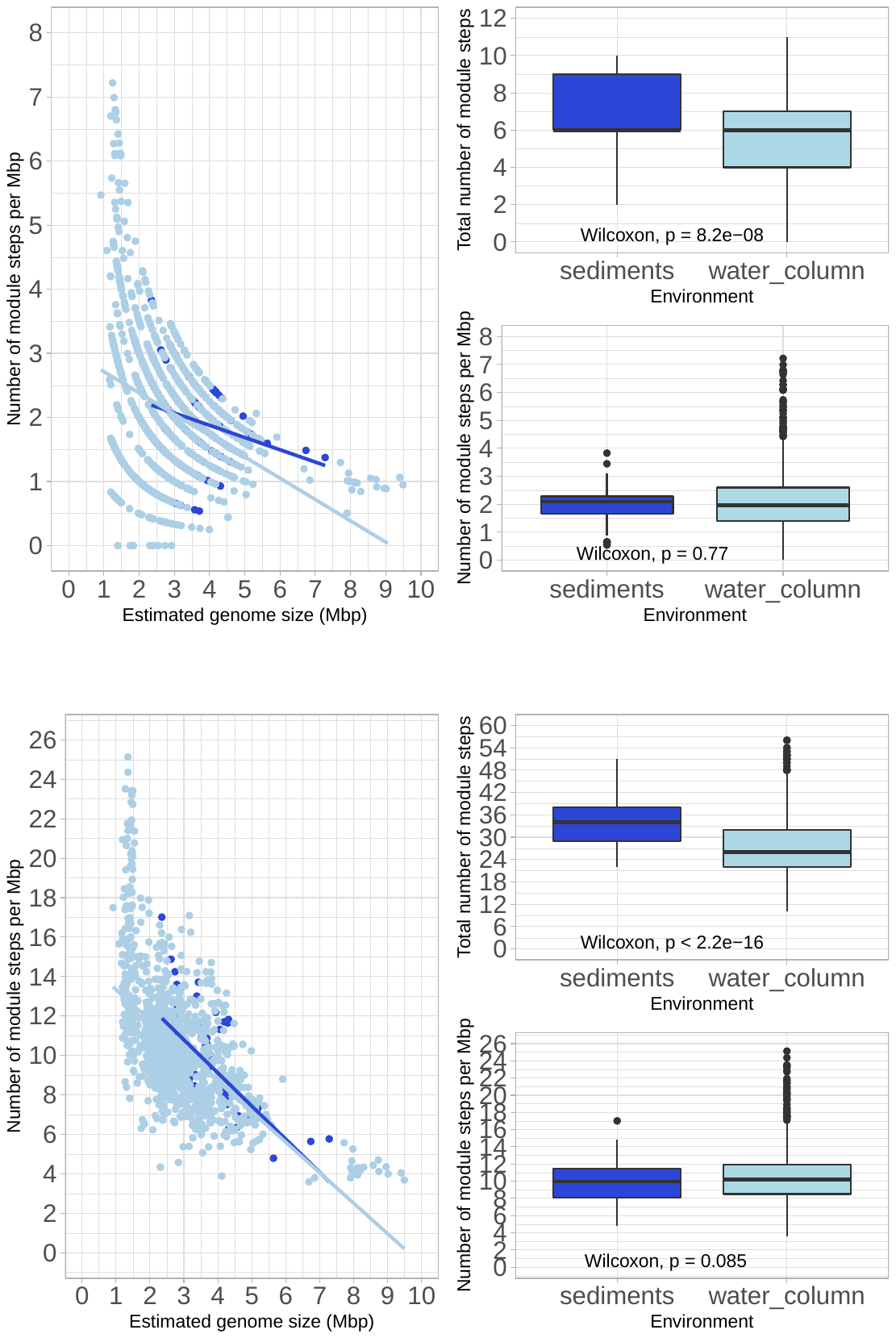
**Nucleotide sugar biosynthesis**

y = 2.6 – 0.19x; R^2^ = 0.067

y = 3 – 0.33x; R^2^ = 0.14

*****

*****

*****

**Other carbohydrate metabolism**

y = 16 – 1.7x; R^2^ = 0.4

y = 15 – 1.5x; R^2^ = 0.36

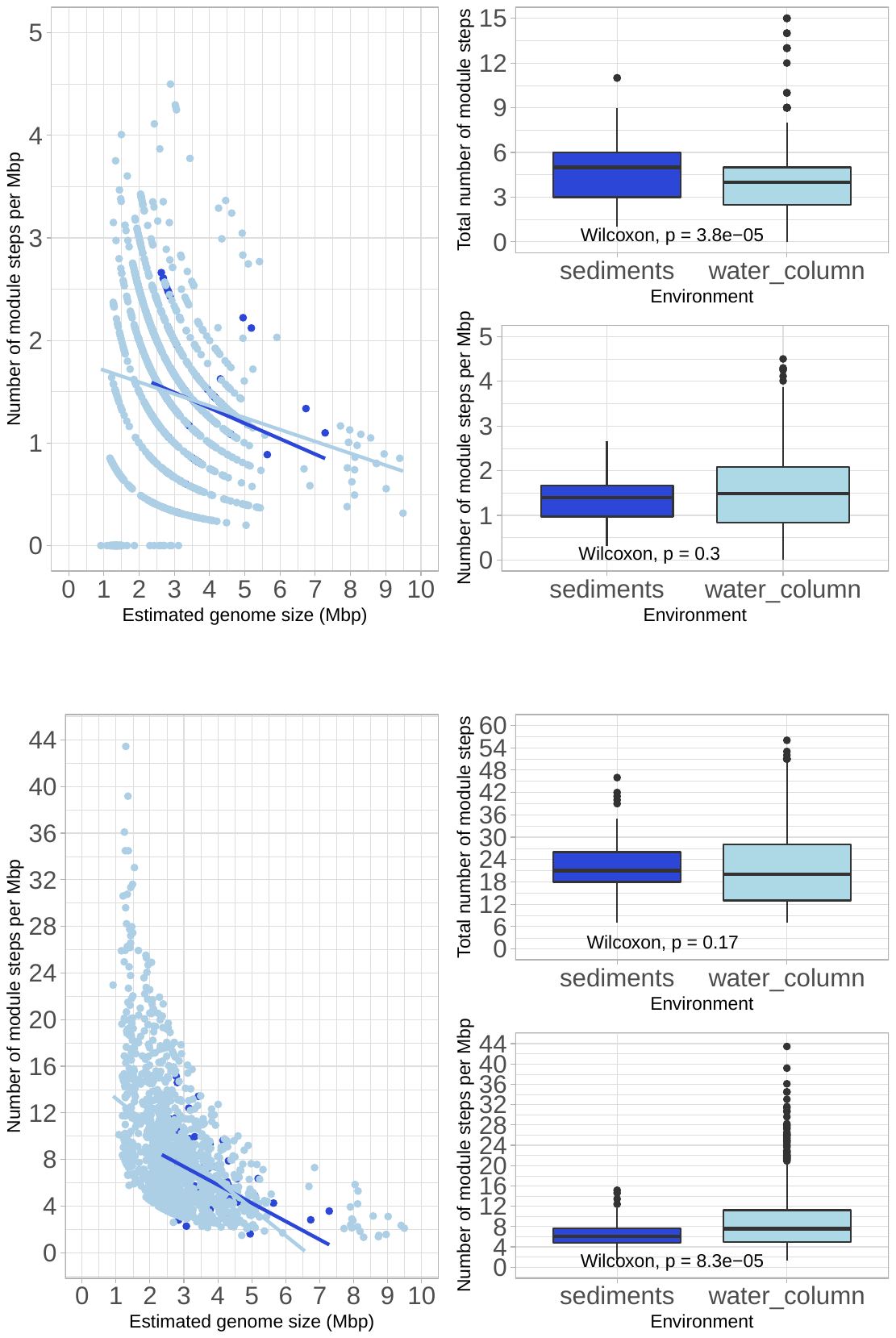
**Polyketide biosynthesis**

y = 1.9 – 0.15x; R^2^ = 0.047

y = 1.8 – 0.12x; R^2^ = 0.027

*****

**Transport systems**

y = 12 – 1.6x; R^2^ = 0.23

y = 16 – 2.4x; R^2^ = 0.26

*****

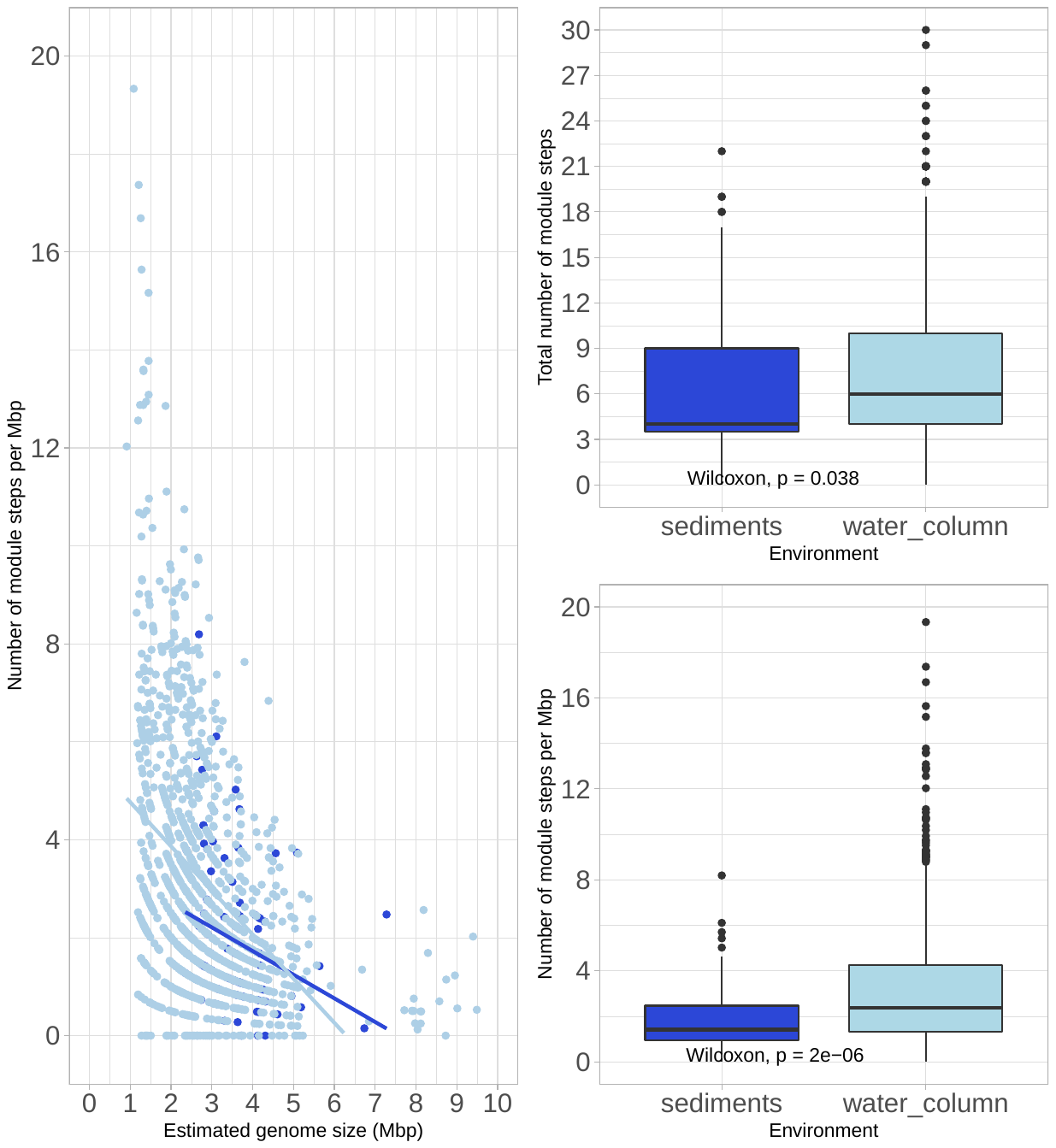
**Two-component regulatory system**

y = 3.7 – 0.48x; R^2^ = 0.072

y = 5.7 – 0.9x; R^2^ = 0.17

*****

*****

List of functional categories used in Figure 3 and Supplementary Figure 2 with the different module categories provided by METABOLIC, and the number of module steps used in the analysis.

Figure 3

Aminoacid metabolism. Figures 3A, 3B and 3C. Module categories:

- Arginine and proline metabolism (21 module steps)
- Lysine metabolism (57 module steps)
- Cysteine and methionine metabolism (35 module steps)
- Serine and threonine metabolism (15 module steps)
- Branched-chain amino acid metabolism (21 module steps)
- Aromatic amino acid metabolism (52 module steps)
- Histidine metabolism (14 module steps)
- Other amino acid metabolism (8 module steps)

Aminoacyl tRNA. Figures 3D, 3E and 3F. Module categories:

- Aminocyl tRNA (42 module steps)

Carbon fixation. Figures 3G, 3H and 3I. Module categories:

- Carbon fixation (108 module steps)

Central carbohydrate metabolism. Figures 3J, 3K and 3L. Module categories:

- Central carbohydrate metabolism (75 module steps)

Nitrogen metabolism. Figures 3M, 3N and 3O. Module categories:

- Nitrogen metabolism (14 module steps)

Sulfur metabolism. Figures 3P, 3Q and 3R. Module categories:

- Sulfur metabolism (7 module steps)

Supplementary Figure 3

Aromatics degradation. Module categories:

- Aromatics degradation (68 module steps)

Beta-lactam and macrolide biosynthesis. Module categories:

- Macrolide biosynthesis (28 module steps)
- Beta-Lactam biosynthesis (22 module steps)

Biosynthesis of secondary metabolites. Module categories:

- Capsaicin biosynthesis (4 module steps)
- Biosynthesis of other secondary metabolites (94 module steps)

Cell signaling. Module categories:

- Cell signaling (75 module steps)

Cofactor and vitamin metabolism. Module categories:

- Cofactor and vitamin metabolism (131 module steps)

Drug resistance. Module categories:

- Drug resistance (115 module steps)
- Drug efflux transporter/pump (54 module steps)

Lipid metabolism. Module categories:

- Lipid metabolism (44 module steps)

Lipopolysaccharides biosynthesis. Module categories:

- Lipopolysaccharide metabolism (18 module steps)
- Lipopolysaccharide biosynthesis, inner core => outer core => O-antigen (12 module steps)

Methane metabolism. Module categories:

- Methane metabolism (52 module steps)

Nucleotide metabolism. Module categories:

- Pyrimidine metabolism (21 module steps)
- Purine metabolism (25 module steps)

Nucleotide sugar biosynthesis. Module categories:

- Nucleotide sugar (12 module steps)

Other carbohydrate metabolism. Module categories:

- Other carbohydrate metabolism (125 module steps)

Polyketide biosynthesis. Module categories:

- Polyketide sugar unit biosynthesis (44 module steps)
- Type II polyketide biosynthesis (38 module steps)

Transport systems. Module categories:

- ABC-2 type and other transport systems (25 module steps)
- Saccharide, polyol, and lipid transport system (42 module steps)
- Phosphate and amino acid transport system (22 module steps)
- Mineral and organic ion transport system (21 module steps)
- Metallic cation, iron-siderophore and vitamin B12 transport system (16 module steps)
- Peptide and nickel transport system (8 module steps)

Two-component regulatory system. Module categories:

- Two-component regulatory system (222 module steps)
